# Prolonged disturbance of proteostasis induces cellular senescence via temporal mitochondrial dysfunction and enhanced mitochondrial biogenesis in human fibroblasts

**DOI:** 10.1101/2020.01.22.916221

**Authors:** Yasuhiro Takenaka, Ikuo Inoue, Takanari Nakano, Masaaki Ikeda, Yoshihiko Kakinuma

## Abstract

Proteolytic activities decline with age, resulting in the accumulation of aggregated proteins in aged organisms. To investigate how disturbance in proteostasis causes cellular senescence, we developed a stress-induced premature senescence (SIPS) model, in which normal human fibroblast MRC-5 cells were treated with either the proteasome inhibitor MG132 or the V-ATPase inhibitor bafilomycin A1 (BAFA1) for 5 days. Time-course studies revealed significant increase in intracellular and mitochondrial reactive oxygen species (ROS) during and after drug treatment. Mitochondrial membrane potential initially decreased, but recovered along with PGC-1α-mediated mitochondrial biogenesis, especially after drug treatment. Mitochondrial antioxidant enzymes SOD2 and GPx4 were temporally depleted in mitochondria on day 1 of the treatment, which in turn could cause excess production of mitochondrial ROS. Extra-mitochondrial SOD2 colocalized with protein aggregates and lysosomes in MG132-treated cells on day 1. SOD2 partially interacted with HSC70 and LAMP2, implying that dysfunctional SOD2 was degraded through chaperon-mediated autophagy (CMA) and caused SOD2 depletion in mitochondria. SIPS induction by MG132 or BAFA1 was partially attenuated by co-treatment with rapamycin, in which generation of excess ROS and mitochondrial biogenesis were suppressed. Rapamycin co-treatment also augmented the upregulation of HSP70 and decreased protein aggregates after drug treatment. Our study proposes a possible pathway from the disturbance of proteostasis to cellular senescence via functional changes in mitochondria.

## 1 INTRODUCTION

Cellular senescence is characterized by permanent cell cycle arrest, morphological changes such as flattened cellular shape, increased levels of lysosomes, upregulation of cyclin-dependent kinase (CDK) inhibitors p16 and p21 (Chang et al., 2000), enhanced activity of the mTOR pathway (Hasty, Sharp, Curiel, & Campisi, 2013), the presence of medium levels of p53 (Leontieva, Gudkov, & Blagosklonny, 2010), and expression of senescence-associated β-galactosidase (SA-β-gal) (Dimri et al., 1995). Normal human fibroblasts cultured *in vitro* have limited proliferative capacity, which is known as the “Hayflick limit”. Limited replicative potential and occurrence of replicative senescence are considered to be caused by shortened telomeres that trigger a DNA damage response (DDR) and eventual irreversible cell cycle arrest. On the other hand, stress-induced premature senescence (SIPS) can be induced by several stressors in young proliferative cells that have long, functional telomeres. Several factors have been reported to serve as inducers of SIPS, including DNA-damaging reagents (Maejima, Adachi, Ito, Hirao, & Isobe, 2008), producers of reactive oxygen species (ROS), expression of oncogenic genes (Moiseeva, Bourdeau, Roux, Deschenes-Simard, & Ferbeyre, 2009), and proteasome inhibitors.

Proteolytic activities and the rate of protein turnover decline in aged animals (Hayashi & Goto, 1998; Ishigami & Goto, 1990) and in humans (Hwang, Hwang, Chang, & Kim, 2007). The decline in proteostasis causes the intracellular accumulation of crosslinked protein aggregates (Reeg & Grune, 2015) and/or autofluorescent materials called lipofuscin (Hohn & Grune, 2013), which are prominent markers of cellular senescence. Previous studies have shown that SIPS is induced in human fibroblasts by treatment with proteasome inhibitors such as MG132. MG132, a tripeptide aldehyde, is a reversible inhibitor of β1 caspase-like, β2 trypsin-like, and β5 chymotrypsin-like activity of the 20S proteasome. At a high dose, MG132 increases ROS levels and GSH depletion in human pulmonary fibroblasts (Park & Kim, 2012), Chinese hamster ovary cells (Maharjan, Oku, Tsuda, Hoseki, & Sakai, 2014), and human leukemic cells (Chao, Chang, Su, & Su, 2014), resulting in cell death. When human fibroblasts, including MRC-5 cells, are treated with MG132 at a low dose for a long period, irreversible growth arrest and premature senescence are induced (Chondrogianni & Gonos, 2004; Chondrogianni et al., 2003; Ukekawa, Maegawa, Mizutani, Fujii, & Ayusawa, 2004). These findings indicate that prolonged disturbance of proteostasis could cause SIPS. However, the exact mechanisms underlying SIPS caused by proteasome inhibitors are unknown.

Mitochondria, the organelles responsible for ATP generation as a source of energy in eukaryotic cells, appear to be the main source of ROS generated in cells. During oxidative phosphorylation (OXPHOS) in mitochondria, electron transfer via an electron transport chain in complexes I and III can cause leakage of electrons (Brand, 2016), which is considered to be the major source of intracellular superoxide, hydrogen peroxide, and other downstream ROS such as hydroxyl radicals. The role of mitochondria in aging is complicated. It has been established that mitochondrial mass increases in fibroblasts that undergo replicative senescence (Lee, Yin, Chi, & Wei, 2002), SIPS (Lee et al., 2002; Passos et al., 2010), and oncogene-induced senescence (OIS)(Moiseeva et al., 2009). In many cases, mitochondria accumulated in senescent cells also show decreased membrane potential, which indicates mitochondrial dysfunction, and produce excess mitochondrial ROS (Korolchuk, Miwa, Carroll, & von Zglinicki, 2017; Nacarelli & Sell, 2017; Passos et al., 2010). Thus, changes in mitochondrial mass and function are the major determinants of cellular senescence.

In this study, we developed a SIPS model by prolonged treatment of young human fibroblast MRC-5 cells, which showed the Hayflick limit at around 60 population doubling level (PDL), with MG132 or bafilomycin A1 (BAFA1) to simulate the situation of proteostasis-impaired cells in aged animals. BAFA1, a plecomacrolide, is a specific and potent inhibitor of vacuolar-type ATPases (V-ATPases), proton pumps that acidify lysosomes, inhibiting lysosomal degradation of biomolecules from several clearance pathways, including macroautophagy and mitophagy (Huss & Wieczorek, 2009).

Both MG132 and BAFA1 impaired cellular proteostasis and induced premature senescence in MRC-5 cells in a similar manner, although they target different molecules. To elucidate the underlying mechanisms of our SIPS model, we performed time-course studies on senescence-associated markers, ROS, DDR, and mitochondrial function and biogenesis during and after treatment with MG132 or BAFA1.

## 2 RESULTS

### 2.1 Prolonged disturbance of proteostasis by MG132 or BAFA1 induces cell cycle arrest, DNA damage response, and SIPS in early passage human fibroblasts

To investigate the mechanism by which prolonged inhibition of proteostasis induces premature senescence in fibroblasts, we first established an MG132- or BAFA1-induced cellular senescence model. We treated normal human fibroblast MRC-5 cells (36-39 PDL) with low concentrations of MG132 or BAFA1 for 3 or 5 days (Fig. 1a). During MG132 or BAFA1 treatment for 5 consecutive days, the cells almost terminated proliferation and acquired thin needle-like morphology (Fig. S1A, day 3). When the culture medium was replaced with the regular medium without the inhibitor (Fig. 1a), sustained growth arrest was observed (Fig. 1b). After drug removal, the cells gradually changed their shape and showed a typical senescence-like morphology, with an enlarged flat shape, on post-treatment day (PD) 5 (Fig. S1A, PD5). However, when cells were treated with the same concentration of MG132 or BAFA1 for 3 consecutive days, proliferation resumed after drug removal (Fig. 1b). Induction of SIPS was also examined using conventional SA-β-gal staining and activity using SPiDER-βGal. Treatment of cells with either of the drugs for 5 days strongly enhanced SA-β-gal-positive cells (Fig. 1c) and SA-β-gal activity (Fig. 1d) after treatment; however, treatment for 3 days did not (Fig. 1c). Therefore, subsequent investigations were focused on cells treated for 5 days with either of the drugs. We also observed a marked increase in the number of aggresome-like inclusion bodies as early as day 1 of treatment with MG132 or BAFA1 (Fig. 1e), which were morphologically similar to those in cells undergoing replicative senescence (58 PDL). Aggregates were detectable even on PD6. DDR induction was examined using immunofluorescence and immunoblotting with anti-activated H2AX (γH2A.X) antibody. DNA damage foci were observed in the nuclei of cells on PD5 (Fig. 1f), and γH2A.X was strongly upregulated in cells after treatment with either of the drugs (Fig. 1g). We next performed time-course analyses of senescence-associated marker proteins. Protein levels of p21 and p53, which are the hallmarks of cellular senescence (Leontieva et al., 2010), were increased as early as day 1, and almost constitutively expressed thereafter, indicating prompt and sustained cell cycle arrest (Fig. 1g). Expression levels of p21 and p53 were comparable to those of aged MRC-5 cells that had undergone replicative senescence (Fig. S1B, 55 and 60 PDL). Sustained activation of mTOR has been reported to be required for cellular senescence (Blagosklonny, 2011; Cho & Hwang, 2012). Phosphorylated ribosomal protein S6 (p-S6) and initiation factor 4E-binding protein 1 (p-4E-BP1), the two best markers of mTOR activity, were depleted once during drug treatment, but recovered to the initial (control) level after the treatment (Fig. 1g and Fig. S2A), indicating restoration of the growth signal in cells that ceased to proliferate after the treatment (Fig. 1b). The reproducibility of protein expression was confirmed with differently prepared samples (Fig. S1C). Both NRF1 and NRF2, NF-E2-related factors 1 and 2, respectively, were transiently upregulated at a very early period of MG132 treatment (Fig. 1h), but diminished thereafter (Fig. S2B). In BAFA1-treated cells, slight induction of NRF1/2 at 48 h was observed (Fig. 1h). We further investigated metabolic remodeling as seen in other cellular senescence models, which showed upregulation of OXPHOS and/or glycolysis (Kaplon et al., 2013; Takebayashi et al., 2015). There seemed to be little increase in glucose transporter GLUT1 and HIF1α protein levels (Fig. S2C), or reduction of PDK1 that inhibits pyruvate dehydrogenase, a key molecule for a respiration (Fig. S2A) in MG132- and BAFA1-treated cells. These findings indicate that there was no metabolic shift in our SIPS model.

**FIGURE 1.**
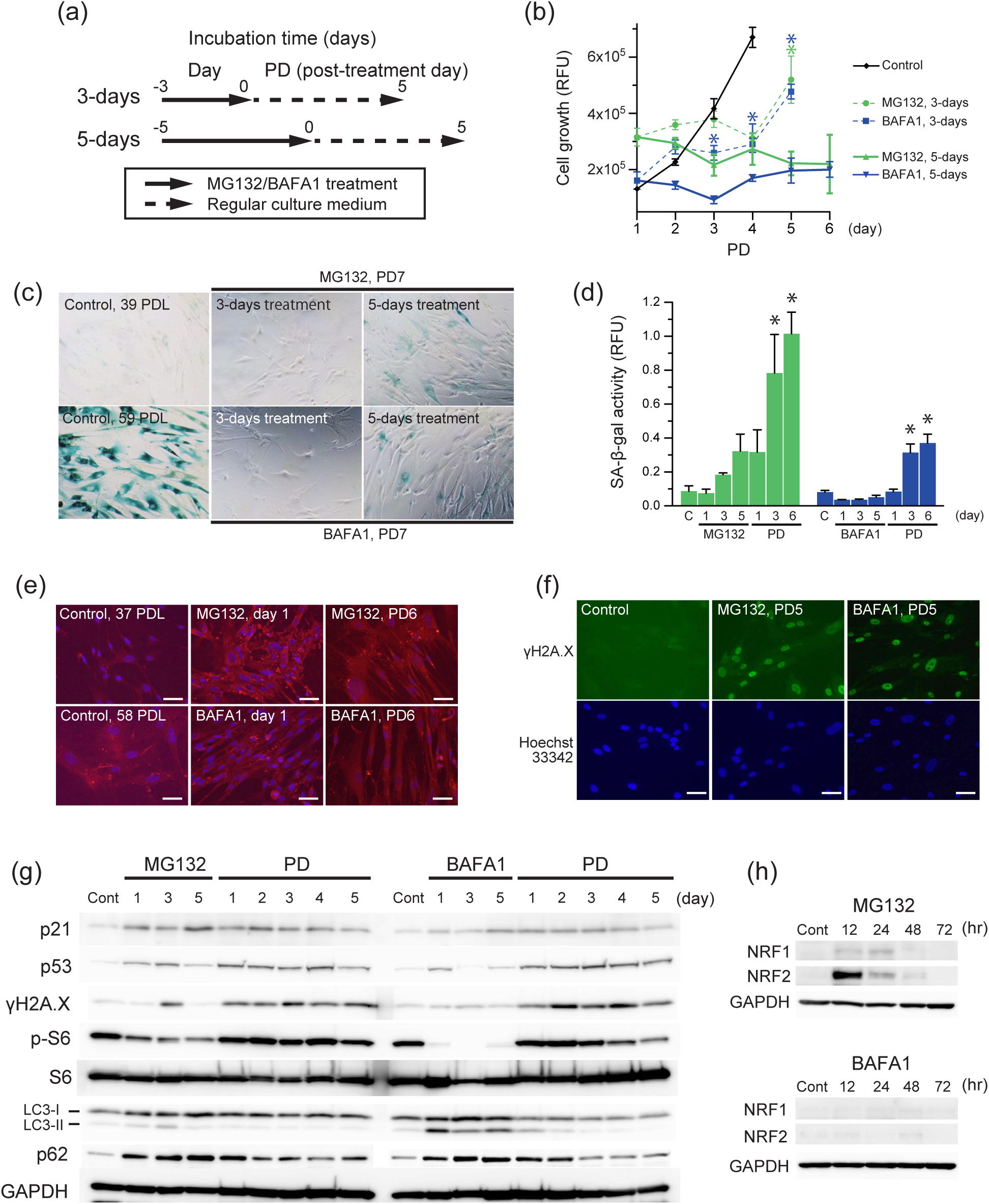
Prolonged disturbance of proteostasis by MG132 or BAFA1 induces premature senescence in normal human fibroblasts. (a) Experimental design for SIPS induced in MRC-5 cells by MG132 or BAFA1. Cells (37-40 PDL) were treated with 0.2-1 μM MG132 or 8-15 nM BAFA1 consecutively for 3 or 5 days without medium refreshment. On day 3 or 5, the drugs were removed and cells were kept in regular culture medium for an additional 5-7 days. Day, day during drug treatment; PD, post-treatment day. (b) Cell proliferation was monitored by measuring Hoechst 33342 fluorescence from PD1 to PD6. Cells were treated with MG132 (green) or BAFA1 (blue) for 3 (dotted lines) or 5 (solid lines) days, and then cultured in medium without the drug for 6 days (PD1-6). Values are shown as the mean (SD) (*n* = 4). The asterisk (*) indicates *p* < 0.05 by one-way ANOVA, followed by Tukey’s multiple comparison test for PD1 vs. others. (c) Representative images of SA-β-gal-positive MRC-5 cells. Cells were treated with MG132 or BAFA1 for 3 or 5 days, respectively, and stained on PD7 (cytoplasmic blue precipitate). Control, untreated young (39 PDL), and old (59 PDL) MRC-5 cells undergoing replicative senescence. (d) SA-β-gal activity was measured using SPiDER-βGal Cellular Senescence Plate Assay Kit. The fluorescence intensity of SA-β-gal activity was normalized to that of Hoechst 33342-stained nuclei. C, untreated control cells. Values are shown as the mean (SD) (*n* = 4). The asterisk (*) indicates *p* < 0.05 by one-way ANOVA, followed by Tukey’s multiple comparison test for control vs. treated cells. (e) Representative images of accumulation of protein aggregates in control and senescence-induced MRC-5 cells. Cells were treated with MG132 or BAFA1 for 5 days, and stained with ProteoStat Aggresome detection reagent (red) and Hoechst 33342 (blue). Scale bar = 50 μm. (f) Representative images of control and senescence-induced MRC-5 cells immunostained for the DNA damage response marker γH2A.X (green) and Hoechst 33342 (blue). Scale bar = 50 μm. (g) Immunoblot analyses of MG132- or BAFA1-treated cells for p21, p53, γH2A.X, downstream target of mTORC1 S6 and phosphorylated S6^Thr235/236^, autophagy marker LC3, and GAPDH. (h) Immunoblot analyses of MG132 (upper)- or BAFA1 (lower)-treated cells for NRF1 and NRF2.

We also tested for SIPS in rat H9c2 myoblasts and mouse NIH3T3 fibroblasts by MG132 or BAFA1 with the same treatment regimen as adopted in MRC-5 cells, and found that senescence induction by these inhibitors is not peculiar to human fibroblasts (Fig. S3A-N).

### 2.2 Prolonged treatment of MG132 or BAFA1 increases intracellular ROS, H_2_O_2_, and mitochondrial ROS, but not nitric oxide

Treatment of cells with various doses (1-30 μM) of MG132 increases intracellular ROS (Legesse-Miller et al., 2012; Park & Kim, 2012) and nitric oxide (NO) production (Chao et al., 2014), but the kinetics of ROS/NO production in fibroblasts during and after longer treatment with MG132 or BAFA1 is unknown. Both intracellular H_2_O_2_ (Figs 2a and 2b) and hydroxyl radicals (.OH) (Figs. S4A and S4B) levels significantly increased from day 3, and were still higher than those of control cells even after drug treatment (PD1 to PD5). In contrast, NO levels remained largely unchanged or slightly reduced during and after treatments (Fig. S4C). There was no increase in mRNA and protein levels of inducible NOS (data not shown), which confirmed that NO was not the cause of senescence induction in MG132- and BAFA1-treated cells. To investigate the involvement of mitochondria in the increased production of intracellular ROS in MG132- or BAFA1-treated cells, we assessed mitochondrial ROS levels using the MitoSOX^TM^ superoxide indicator. Mitochondrial ROS levels significantly increased in both MG132- and BAFA1-treated cells as early as day 1 of the treatment, and peaked at PD3 (Figs. 2c and 2d).

**FIGURE 2.**
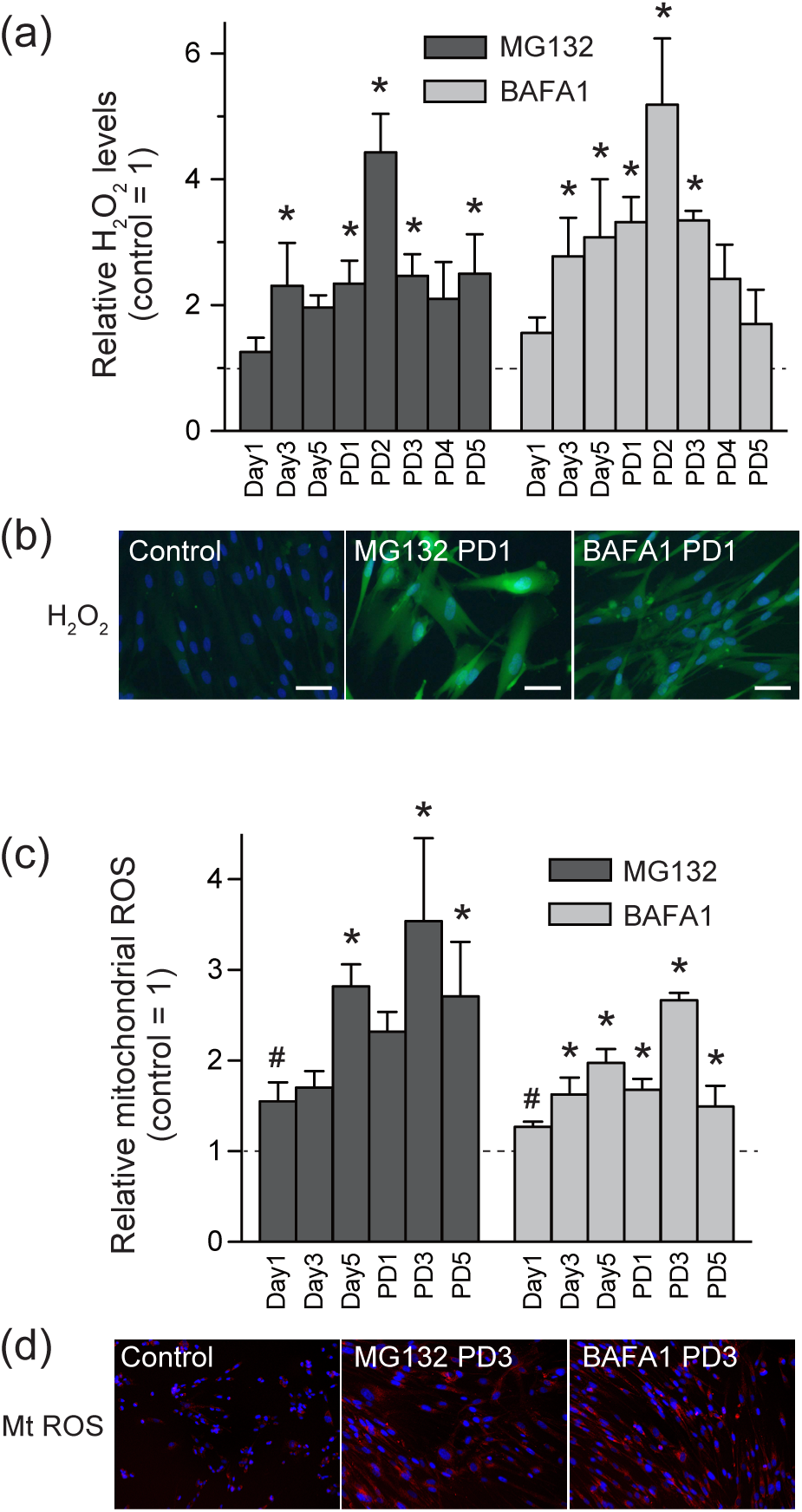
Prolonged disturbance of proteostasis by MG132 or BAFA1 enhances ROS production. (a) Quantitation of relative intracellular H_2_O_2_ levels using HYDROP reagent that selectively detects H_2_O_2_. The fluorescence intensity for HYDROP was normalized by total protein concentration. Values are shown as the mean (SD) (*n* = 4). (b) Representative images of control and MG132- or BAFA1-treated cells stained with HYDROP (green) and Hoechst 33342 (blue) on PD1. Scale bar = 50 μm. (c) Relative mitochondrial ROS levels were examined using MitoSOX Red mitochondrial superoxide indicator. The fluorescence intensities for MitoSOX and Hoechst 33342 were quantified by ImageJ. The fluorescence intensity for MitoSOX was normalized by the number of nuclei. Values are shown as the mean (SD) from the measurements of four different images. The asterisk (*) indicates *p* < 0.05 by one-way ANOVA, followed by Tukey’s multiple comparison test for control vs. treated cells. Values obtained from cells treated with MG132 or BAFA1 on day 1 were compared with those from untreated control cells using a non-parametric Mann-Whitney *U* test. The sharp (#) indicates significant difference (*p* < 0.05). (d) Representative images of control and MG132- or BAFA1-treated cells stained with MitoSOX (red) and Hoechst 33342 (blue) on PD3.

### 2.3 Prolonged treatment of MG132 or BAFA1 enhances mitochondrial biogenesis

We next investigated changes in mitochondrial mass and mitochondrial membrane potential (Δψm) using MitoTracker Green FM and MitoTracker Red CM-H2Xros, respectively. In cells treated with either drug, the mitochondrial mass gradually increased during treatment and remained at high levels even after treatment (Fig. 3a and Mt mass in Fig. 3c). In contrast, the mitochondrial membrane potential (Δψm) decreased slightly on days 1, 3, and 5, compared with untreated control cells, and markedly increased thereafter (Figs. 3b, 3c, and S5E). Mitochondrial proliferation was also confirmed by qPCR analysis of mitochondrial DNA (mtDNA) copy number (Fig. 3d). Intracellular ATP levels were almost at basal levels during treatment, but significantly increased thereafter (Fig. 3e), possibly reflecting the restoration of mitochondrial membrane potential after drug treatment. These results suggest that there was temporal mitochondrial dysfunction during drug treatment, but an increase in functional mitochondria, probably by mitochondrial biogenesis thereafter.

**FIGURE 3.**
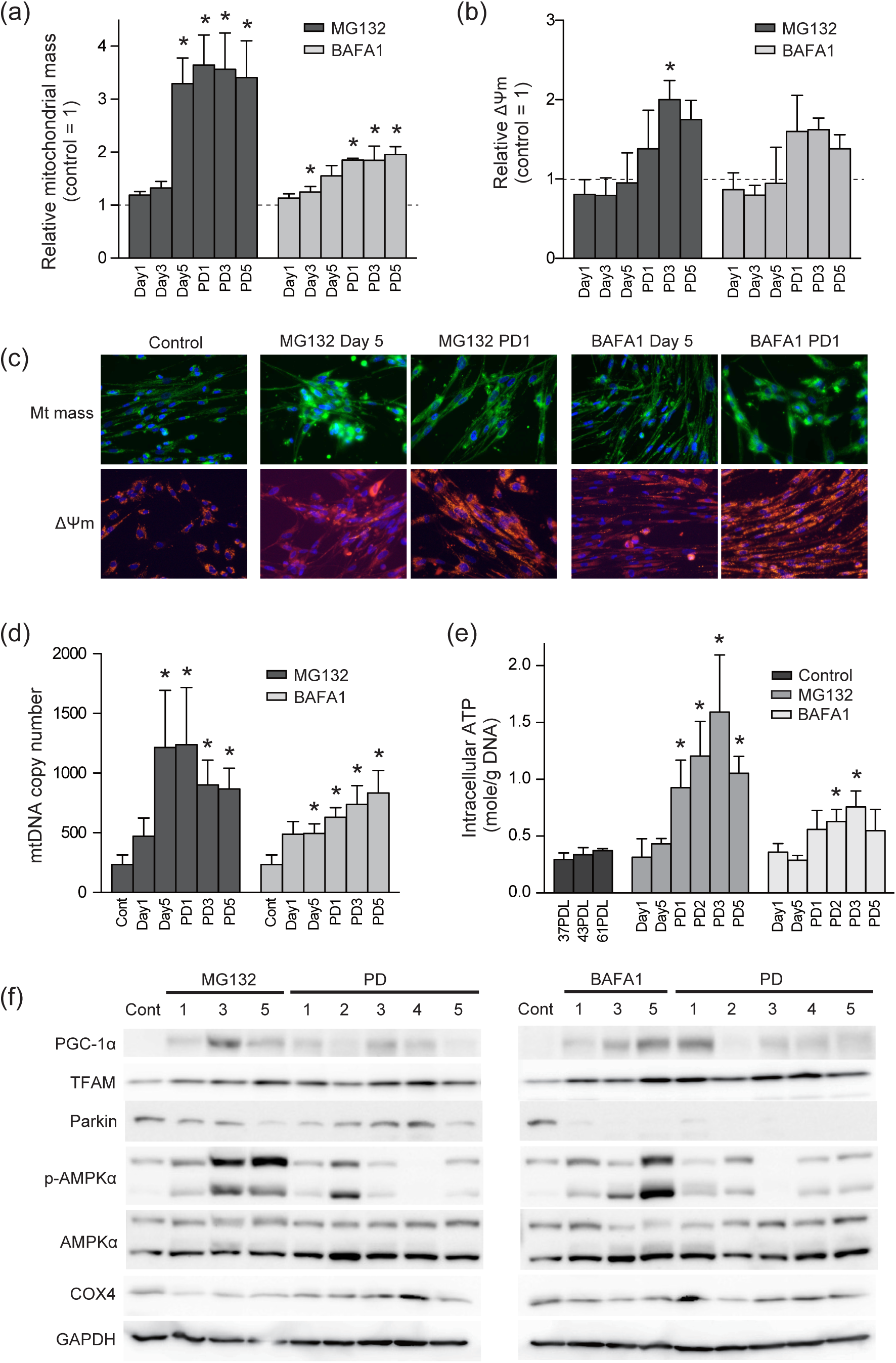
Prolonged disturbance of proteostasis by MG132 or BAFA1 induces temporal mitochondrial dysfunction and sequential mitochondrial biogenesis. (a) Relative mitochondrial mass was evaluated using MitoTracker Green FM. The fluorescence intensities for MitoTracker Green and Hoechst 33342 were quantified using ImageJ. MitoTracker Green intensity was normalized by the number of nuclei. Values are shown as the mean (SD) from the measurements of five different images. The asterisk (*) indicates *p* < 0.05 by one-way ANOVA, followed by Tukey’s multiple comparison test for control vs. treated cells. Relative mitochondrial membrane potential (ΔΨm) was evaluated using MitoTracker Red CM-H_2_XRos. The fluorescence intensities for MitoTracker Red CM-H_2_XRos and Hoechst 33342 were quantified using ImageJ. MitoTracker Red CM-H_2_XRos intensity was normalized by the number of nuclei. Values are shown as the mean (SD) from the measurements of 10 different images. The asterisk (*) indicates *p* < 0.05 by one-way ANOVA followed by Tukey’s multiple comparison test for control vs. treated cells. (c) Representative images of control and MG132- or BAFA1-treated cells stained with MitoTracker Green FM (Mt mass, green), MitoTracker Red CM-H_2_XRos (ΔΨm, Red) and Hoechst 33342 (blue) on day 5 and PD1. (d) The copy number of mitochondrial DNA (mtDNA) of control and MG132- or BAFA1-treated cells was measured by qPCR using mtDNA-specific primers and nuclear DNA-specific primers as a reference. Values are the mean (SD) (*n* = 4). The asterisk (*) indicates *p* < 0.05 by one-way ANOVA, followed by Tukey’s multiple comparison test for control vs. treated cells. (e) Intracellular ATP contents were measured using luciferase-based assay. The luminescence of ATP was normalized by cellular DNA concentration quantified using SYBR Gold. Values are shown as the mean (SD) (*n* = 4). The asterisk (*) indicates *p* < 0.05 by one-way ANOVA, followed by Tukey’s multiple comparison test for control (37 PDL) vs. treated cells. (f) Immunoblot analyses of control and MG132 (left)- or BAFA1 (right)-treated MRC-5 cells for PGC-1α, TFAM, Parkin, AMPKα, phosphorylated AMPKα^Thr172^ (p-AMPKα), COX4 and GAPDH.

We further explored how mitochondrial mass increased in MG132- or BAFA1-treated cells by assessing the protein levels of key factors in mitochondrial biogenesis and clearance. PGC-1α, the master regulator of mitochondrial biogenesis, was upregulated temporarily from day 3 to PD1 with different kinetics in MG132 and BAFA1, but was downregulated thereafter (Fig. 3f). In contrast, TFAM, the key mitochondrial transcription activator, was consistently elevated during and after treatment. Upregulation of these factors indicated enhanced mitochondrial biogenesis in MG132- and BAFA1-treated cells. Parkin, the ubiquitin ligase that is responsible for mitophagy, was largely downregulated below basal levels, indicating suppression of mitophagic flux during and after treatment.

We also found that AMP-activated protein kinase alpha (AMPKα) was strongly activated, especially on days 3 and 5 (Fig. 3f), which was consistent with the increase in intracellular ATP levels in the post-treatment periods (Fig. 3e). Increased cytochrome c oxidase subunit 4 (COX4) levels also confirmed an increase in mitochondrial mass in MG132- and BAFA1-treated cells.

### 2.4 MG132- or BAFA1-treatment causes reduction in mitochondrial antioxidant proteins in mitochondrial fraction during the early period of treatment

Recently, mass spectrometry analysis revealed aggregation of nuclear-encoded mitochondrial proteins such as respiratory chain complex subunits to be an early event in MG132-treated cells (Rawat, Anusha, Jha, Sreedurgalakshmi, & Raychaudhuri, 2019). In our experiments, mitochondrial ROS was significantly increased in MG132- or BAFA1-treated cells as early as day 1, compared with untreated control cells (Fig. 2c), implying that enhanced ROS generation and/or failure to scavenge ROS occurred in the mitochondria immediately after initiation of drug treatment. Thus, we investigated the protein levels of two mitochondrial antioxidant enzymes, superoxide dismutase (SOD2) and glutathione peroxidase (GPx4), in the isolated mitochondrial fraction (detailed method in Fig. S6A-C) of MG132- or BAFA1-treated cells. On day 1, SOD2 was significantly depleted in the mitochondrial fraction (Mt) (Figs. 4a-d and S7E) in cells treated with either drug cells although the overall SOD2 level (whole) was almost constant compared with control cells (Figs. 4a, 4b, S7A, and S7C). From day 5, whole and mitochondrial SOD2 levels increased markedly. On the other hand, GPx4 protein levels were significantly depleted not only in Mt but also in the whole fraction on day 1. The protein levels of TOM40 and GAPDH in the mitochondrial fraction remained largely unchanged in cells treated with either drug. To further clarify the cause of GPx4 protein reduction in the whole fraction, we examined *GPx4* mRNA levels by qPCR, and found a significant reduction in the amount of transcripts in MG132-treated cells (Fig. S7B) and a slight reduction in BAFA1-treated cells on day 1 (Fig. S7D). Thus, GPx4 depletion in Mt and whole fraction possibly arose from the reduction of GPx4 transcripts. We next investigated SOD2 localization to elucidate the mechanism of the protein depletion in the Mt fraction on day 1. We observed that SOD2 partially accumulated in the area of extra-mitochondrial (COX4 negative) regions in both MG132- and BAFA1-treated cells on day 1, whereas SOD2 largely colocalized with COX4 in control cells (Figs. 4e and S8A). We also found that the extra-mitochondrial SOD2 signals partially colocalized with aggregated protein signals (Figs. 5a, 5b, S9A, and S9B), suggesting that SOD2 could form aggregates before translocation into mitochondria. To investigate the formation of SOD2 aggregates, we analyzed detergent-insoluble fractions (pellets of 0.5% Triton X-100 lysate) of MG132- or BAFA1-treated cells, but found that there was no increase in insoluble SOD2 by drug treatment (data not shown). Then, we fixed MG132- or BAFA1-treated cells with paraformaldehyde, prepared detergent-soluble cellular lysates, and examined the association of SOD2 with other proteins by immunoblotting. We found that signals of high molecular mass SOD2 significantly increased in MG132- and BAFA1-treated cells than in control cells (Fig. 5d), thereby suggesting enhanced complex formation of SOD2 with other proteins in drug-treated cells. When the lysates were heat-denatured at 95°C for 5 min to partly cleave crosslinks, the monomeric and multimeric SOD2, but no SOD2 complexes as seen in non-heated samples, appeared in all samples. GAPDH was largely monomeric, even upon fixation. Next, we explored the modification and interactions of SOD2 with other proteins. We detected neither ubiquitination nor acetylation of SOD2 on day 1 (Fig. S8B and data not shown), but we found partial interaction of SOD2 with chaperon protein HSC70 and lysosome membrane protein LAMP2 (Fig. 5e), implying lysosomal degradation of SOD2 by chaperon-mediated autophagy (CMA). Protein sequence analysis using a KFERQ finder (Kirchner et al., 2019) revealed two putative KFERQ-like motifs, which can be targeted by HSC70 (_128_FDKFK_132_ and _130_KFKEK_134_) in the SOD2 polypeptide sequence. In accordance with these results, SOD2 partially colocalized with lysosomes (LAMP1-GFP) in MG132-treated cells on day 1 (Figs. 5f and S9D).

**FIGURE 4.**
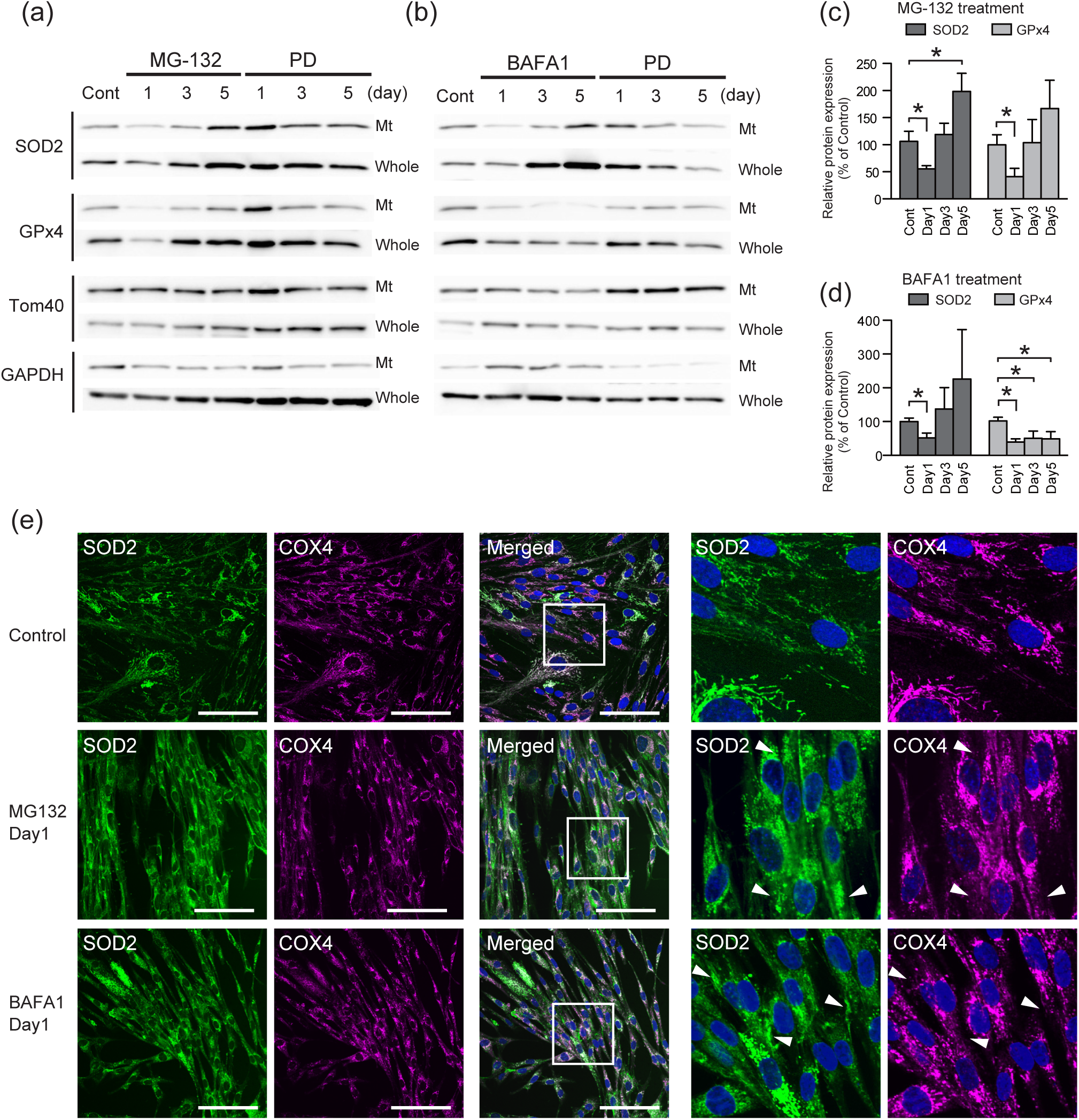
Depletion of the mitochondrial antioxidant enzymes SOD2 and GPx4 in mitochondrial fraction in the early period of MG132- or BAFA1-treated MRC-5 cells. (a and b) Immunoblot analyses of MG132 (a)- or BAFA1 (b)-treated cells for SOD2, GPx4, TOM40 (mitochondrial loading control), and GAPDH (cytosolic loading control). Mt, crude mitochondrial fraction; Whole, whole cellular extract; Cont, control cells. Five (mitochondrial fraction) or 15 (whole cellular extract) μg of proteins were loaded onto each lane. (c and d) Quantitative analyses of the mitochondrial SOD2 and GPx4 protein levels in MG132 (b)- or BAFA1 (d)-treated cells. Intensities of SOD2 and GPx4 bands were normalized to that of GAPDH. Values are shown as the mean (SD) (*n* = 4). The asterisk (*) indicates *p* < 0.05 by a non-parametric Mann-Whitney *U* test for control vs. treated cells. (e) Representative images of control and MG132- or BAFA1-treated MRC-5 cells immunostained for SOD2 (green), COX4 (purple), and Hoechst 33342 (blue) on day 1. Arrowheads indicate extra-mitochondrially localized SOD2. Scale bar = 100 μm.

**FIGURE 5.**
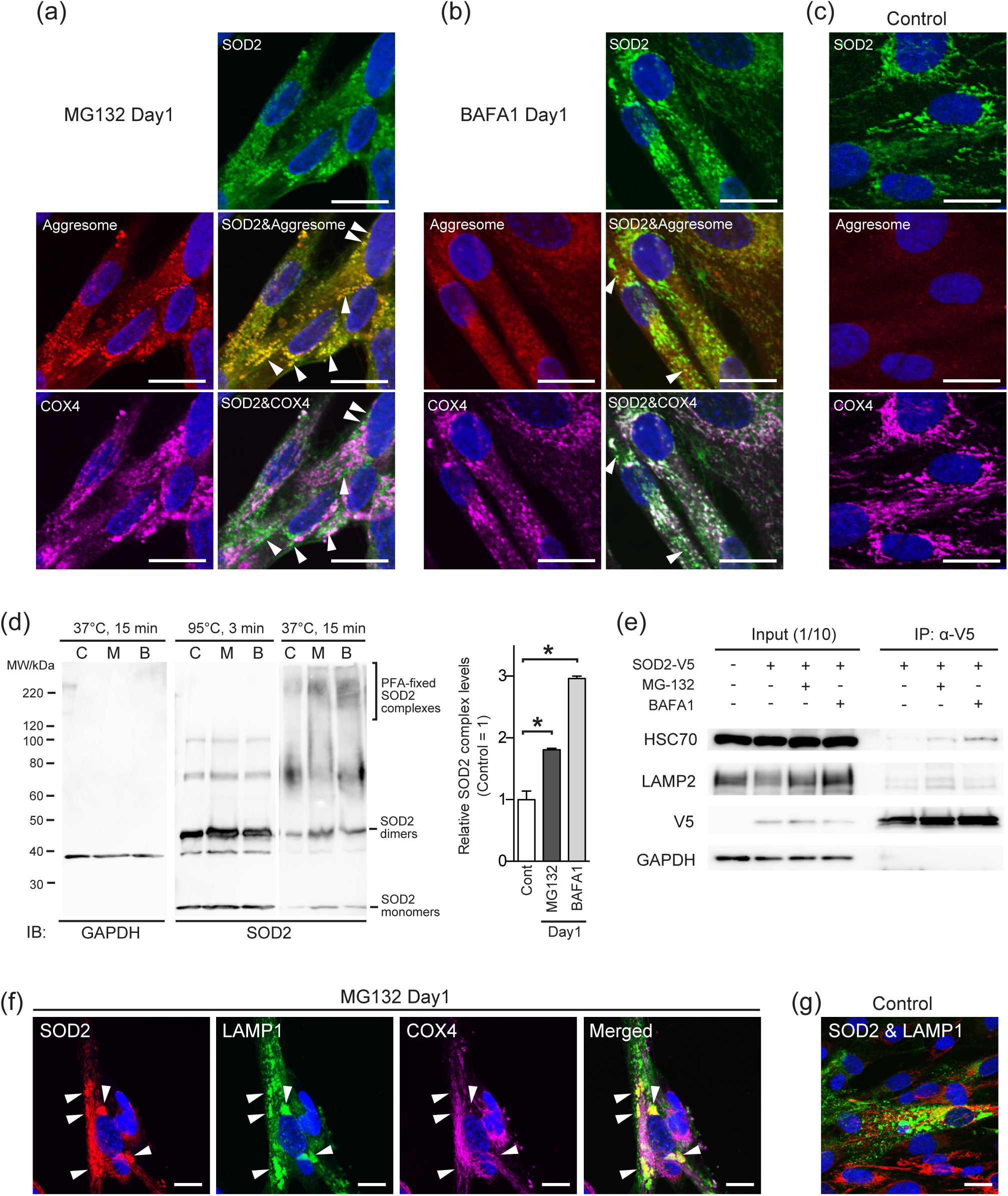
Extra-mitochondrial SOD2 colocalizes with aggregated proteins and lysosome. (a and b) Representative images of MRC-5 cells treated with MG132 (a) or BAFA1 (b) for a day and immunostained for SOD2 (green) and COX4 (purple). Cells were further stained using the protein-aggregate detection reagent (ProteoStat, red) and Hoechst 33342 (blue). Arrowheads indicate extra-mitochondrial SOD2 that colocalized with aggresome. (c) Immunostained and ProteoStat-stained control cells. Scale bar = 20 μm. Immunoblot analyses of PFA-fixed SOD2 complexes in cells treated with MG132 or BAFA1 on day 1. Left panels, lysates prepared from PFA-fixed cells were incubated at 37°C for 15 min or 95°C for 3 min, loaded onto SDS-PAGE (5 μg∕lane), and immunoblotted (IB) with anti-SOD2 or GAPDH antibodies. C, M, and B indicate control, MG132-, and BAFA1-treated cells, respectively. Right graph, quantitative analyses of PFA-fixed SOD2 complex levels in MG132- or BAFA1-treated cells on day 1. Intensity of SOD2 complexes was normalized to that of GAPDH. Values are shown as the mean (SD) (*n* = 4). The asterisk (*) indicates *p* < 0.05. (e) Direct interaction of SOD2-V5 and HSC70 in MG132- or BAFA1-treated cells on day 1. (f) Extra-mitochondrial SOD2 (red) colocalizes with lysosomal protein LAMP1 (green) in MG132-treated MRC-5 cells on day 1. LAMP1-GFP fusion protein was overexpressed by CellLight^TM^ Lysosomes-GFP BacMam 2.0. Arrowheads indicate extra-mitochondrial SOD2 that colocalized with LAMP1. (g) Differential localization of SOD2 and LAMP1 in control cells. Scale bar = 20 μm.

### 2.5 Rapamycin treatment partially attenuates cellular senescence induced by MG132 or BAFA1

Mitochondrial function and biogenesis are controlled by mTOR (Morita et al., 2015). Inhibition of mTOR with rapamycin, an mTORC1 inhibitor, resulted in a decrease in mitochondrial oxidative function (Ramanathan & Schreiber, 2009), mitochondrial ROS (Martinez-Cisuelo et al., 2016), and senescence induction (Leontieva, Demidenko, Gudkov, & Blagosklonny, 2011; Lerner et al., 2013; Summer et al., 2019). To further investigate the involvement of mitochondrial biogenesis in our SIPS model, we treated cells with rapamycin and either MG132 or BAFA1. Following 5 days co-treatment with rapamycin and MG132 or BAFA1 from day 0 (Fig. 6a, Rapa1), the cells recovered proliferation potential after PD5 (Fig. 6b). In contrast, poor growth was observed in cells co-treated for 2 days from day 3 and additionally treated with rapamycin for 3 days from PD1 (Fig. 6a and 6b, Rapa2). The effects of rapamycin on SIPS were also confirmed by SA-β-gal activity (Fig. 6c and S10) and intracellular H_2_O_2_ levels (Fig. 6d and 6e) in both MG132- and BAFA1-treated cells. An increase in mitochondrial mass was significantly repressed by co-treatment with rapamycin and MG132 (Rapa1) or BAFA1 (Rapa1 and Rapa2) (Fig. 6f). Different profiles of proliferative potential, H_2_O_2_ levels, and mitochondrial mass between Rapa1 and Rapa2 suggested that cellular events during the early period (days 0-3) of drug treatment strongly affected SIPS progression. Thus, we further analyzed the effect of rapamycin on ROS production during the early period (days 1-2) of the treatment. Mitochondrial ROS was significantly decreased by co-treatment with rapamycin on day 1 (MG132) or day 2 (BAFA1) compared with cells treated with either MG132 or BAFA1 (Fig. 6g). We also found that intracellular aggregation was precipitated by co-treatment with rapamycin and MG132 or BAFA1 on PD3 (Fig. 6h), implying rapamycin treatment improved proteostasis. Rapamycin co-treatment augmented the upregulation of chaperon protein HSP70 in MG132- or BAFA1-treated cells on PD1 (Figs. 6i-k) and the total content of ubiquitinated proteins in MG132-treated cells on day3 (Fig. S11A). To explore how rapamycin promoted cell proliferation (Fig. 6b), we investigated protein levels of the phosphorylated Akt at Ser473 (S473) and Thr308 (T308). S473 level was significantly increased by Rapa1 co-treatment on PD3 (Figs. S11A-D), and changes in T308 largely paralleled those observed on S473. Akt activation was not observed in control cells treated with rapamycin alone for 5 days (Fig. S11E). There was no significant change in p62 and HSC70 protein levels between single treatment with MG132 or BAFA1 and co-treatment with rapamycin (Figs. S11A, S11B, and data not shown). The levels of the active form of LC3 (LC3-II) were not altered by co-treatment with rapamycin compared to cells without rapamycin (Fig. S12), suggesting that autophagic flux for damaged proteins or mitochondrial clearance did not contribute to the rapamycin effect in our SIPS models.

**FIGURE 6.**
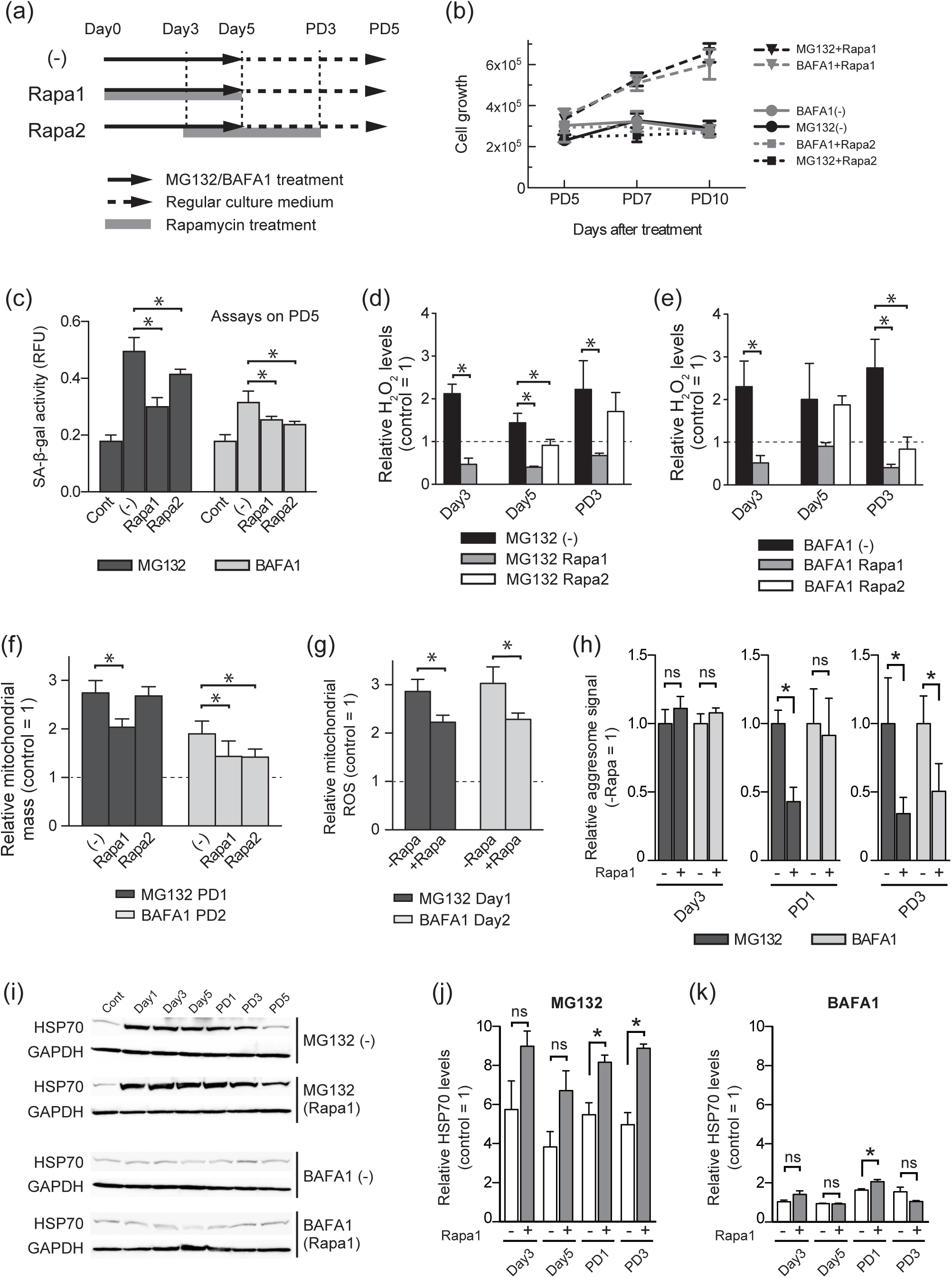
Rapamycin attenuates cellular senescence induced by MG132 or BAFA1. (a) The experimental design for co-treatment of rapamycin (final 10 nM) with MG132 or BAFA1 in MRC-5 cells. Rapa1, cells were co-treated with rapamycin and either MG132 or BAFA1 from day 0 to day 5. Rapa2, cells were co-treated with rapamycin and either MG132 or BAFA1 from day 3 to day 5, and continuously cultured in regular medium containing rapamycin from day 5 to PD3. (b) Cell proliferation was evaluated by measuring Hoechst 33342 fluorescence from PD5 to PD10. Values are shown as the mean (SD) (*n* = 4) (c) SA-β-gal activities of cells co-treated with rapamycin and either MG132 or BAFA1 were measured on PD5 using a SPiDER-βGal Cellular Senescence Plate Assay Kit. Normalized values are shown as the mean (SD) (*n* = 4). Values obtained from cells co-treated with rapamycin and either MG132 or BAFA1 (Rapa1 and Rapa2) were compared with those from cells treated with only MG132 or BAFA1 (-) using a non-parametric Mann-Whitney *U* test (\**p* < 0.05). (d and e) Intracellular H_2_O_2_ levels of cells co-treated with rapamycin and either MG132 or BAFA1 were measured on day 3, day 5, and PD3 by HYDROP staining. Normalized values are shown as the mean (SD) (*n* = 4). Two groups were compared using a non-parametric Mann-Whitney *U* test (\**p* < 0.05). (f) Relative mitochondrial mass of cells co-treated with rapamycin and either of MG132 or BAFA1 was measured on PD1 (MG132) and PD2 (BAFA1) using MitoTracker Green FM. Values are shown as the mean (SD) from the measurements of four different images. Two groups were compared using a non-parametric Mann-Whitney *U* test (\**p* < 0.05). (g) Relative mitochondrial ROS levels of cells co-treated with rapamycin (+Rapa) and either of MG132 or BAFA1 were measured on day 1 (MG132) and day 2 (BAFA1). The MitoSOX intensity was normalized by the number of nuclei. Values are shown as the mean (SD) from the measurements of 4 different images. Two groups were compared using a non-parametric Mann-Whitney *U* test (\**p* < 0.05). (h) Relative protein aggregation levels in cells treated with MG132 or BAFA1 (-Rapa) or co-treated with rapamycin and either MG132 or BAFA1 (+Rapa1) on day 3, PD1, and PD3. Normalized values are shown as the mean (SD) (*n* = 7). Two groups were compared using a non-parametric Mann-Whitney *U* test (\**p* < 0.05). (i) Immunoblot analyses of MG132- or BAFA1-treated cells with (Rapa1) or without (-) rapamycin using anti-HSP70 antibody. (j and k) Quantitative analyses of the HSP70 protein levels in MG132 (j)- or BAFA1 (k)-treated cells. HSP70 protein levels were evaluated by densitometry of each band obtained by immunoblot analyses of four individually prepared whole cellular lysates. Values are shown as the mean (SD). Two groups were compared using a non-parametric Mann-Whitney *U* test (\**p* < 0.05). ns, not significant.

## 3 DISCUSSION

Most previous studies on cellular senescence have focused on changes in metabolism, morphology, DNA damage, and senescence-related gene expression before and after senescence. However, we cannot claim to have elucidated the mechanism of senescence unless we thoroughly understand the process of cellular senescence from initiation to the irreversible state of senescence. Although this work may be a considerable challenge in the case of organismal aging, it is relatively straightforward in the case of cellular senescence, especially SIPS. In the present study, we performed the time-course study of cellular senescence triggered by prolonged inhibition of proteostasis. We found that there are two phases of mitochondrial status in which mitochondrial membrane potential only temporarily decreased during MG132 or BAFA1 treatment, but significantly increased thereafter (Figs. 3b and S5E), along with a significant increase in mitochondrial mass (Fig. 3a and 3d) and ATP levels (Fig. 3e), suggesting that an increased functional mitochondria during the latter period of SIPS progression is one of the major determinants of excess ROS production and eventual DDR. Enhanced mitochondrial biogenesis was confirmed by a notable increase in PGC-1α and TFAM proteins (Fig. 3f), which are critically involved in mitochondrial biogenesis, whereas Parkin (Fig. 3f) and LC3-II (Fig. 1g) were greatly downregulated, implying that mitophagic flux was suppressed during enhanced mitochondrial biogenesis (day 5 to PD2).

We observed a marked depletion of the antioxidant enzymes SOD2 and GPx4 in the mitochondrial fraction (Figs. 4a-d) in MG132- or BAFA1-treated cells on day 1. Extra-mitochondrial SOD2 also showed partial colocalization with aggresome-like inclusions (Figs. 4e, 5a-c) and lysosomes (Fig. 5f) in MG132-treated cells on day 1, implying that cytosolic SOD2 was insufficiently translocated into the mitochondria, as an early event of disturbance of proteostasis. However, in case of BAFA1 treatment, the occurrence of colocalization of SOD2 with lysosomes was lower than that of MG132 (data not shown), and further analysis is required. The association between SOD2 and HSC70 suggested that the extra-mitochondrial SOD2 was degraded by CMA (Fig. 5e). Thus, it is likely that unfolded or damaged SOD2 accumulated in the cytosol by impaired proteostasis, which caused depletion of mitochondrial SOD2 content, and eventually degraded by CMA. On the other hand, depletion of mitochondrial GPx4 protein levels was possibly due to the reduction in the overall protein and transcription levels (Figs. S7A-D) although the mechanism of transcriptional regulation remains to be elucidated.

Unexpectedly, SIPS induced by MG132 and BAFA1 showed similar phenotypes and kinetics of senescence-associated markers, mitochondrial dysfunction and biogenesis, ROS levels, and DDR, although these reagents inhibit different molecules. This observation implies the presence of shared mechanisms in SIPS induced by disturbance of proteostasis, regardless of the initial trigger. In contrast, there were several differences between the profiles of SIPS by MG132 and BAFA1. For instance, the levels of SA-β-gal activity, NRF1/2 induction, and mitochondrial biogenesis were relatively low in BAFA1-treated cells. In addition, the overall progression of SIPS in BAFA1-treated cells seemed to be slower than in MG132-treated cells. Therefore, we could not exclude the presence of drug-specific induction mechanisms in our SIPS model.

mTOR inhibition using rapamycin partially attenuated SIPS induced by MG132 or BAFA1 (Figs. 6b and 6c) through suppression of intracellular H_2_O_2_ (Figs. 6d and 6e), mitochondrial ROS production (Fig. 6g), and mitochondrial biogenesis (Fig. 6f), compared with cells treated with MG132 or BAFA1. In addition, we found that rapamycin co-treatment partially restored proteostasis after drug treatment (Fig. 6h). Both the heat shock protein chaperone system and the ubiquitin-proteasome system might be involved in the effect of rapamycin on proteostasis in our model. Rapa1-treatment enhanced the levels of the chaperone HSP70, especially in MG132-treated cells (Figs. 6i-k). To the best of our knowledge, this is the first report of the effect of rapamycin on HSP70 expression, although the induction mechanism remains unknown. Several reports have shown that rapamycin-mediated inhibition of mTORC1 increases the level of ubiquitinated proteins *in vivo* and *in vitro* (Harston et al., 2011; Zhao, Zhai, Gygi, & Goldberg, 2015). In our model, rapa1-treatment markedly increased the total content of ubiquitinated proteins in MG132-treated cells on day 3 (Fig. S11A). This is a paradoxical response, because prolonged accumulation of ubiquitinated proteins seems to be deleterious to proteasome-inhibited cells. However, proteasomal activity impaired on day 1 gradually recovered, even during MG132 treatment, and returned to control levels on PD1 (data not shown). Therefore, enhanced protein ubiquitination by rapamycin co-treatment would benefit proteostasis, and attenuate the subsequent progression of senescence. Another possibility is that different types of ubiquitin chains are formed on day 3 than on day1. While K48-linked polyubiquitination leads to degradation of the target proteins to 26S proteasomes, K63-linked chains do not (Nathan, Kim, Ting, Gygi, & Goldberg, 2013). In our analysis, we examined the total content of ubiquitinated proteins using anti-Ub antibody. Therefore, the significance of rapamycin-induced ubiquitination in the senescence process awaits further investigation. We also observed Akt activation in MG132- and BAFA1-treated cells by the rapamycin co-treatments (Figs. S11A-D), which suggested contribution of Akt to cell survival and proliferation after drug treatment. Although rapamycin-mediated Akt activation has been reported previously (Dibble, Asara, & Manning, 2009; Harston et al., 2011), it is noteworthy that phosphorylated Akt levels mainly increased in the post-treatment period (PD1-5) in an MG132- or BAFA1-dependent manner. Nogueira et al. reported that Akt1/2 null primary mouse embryonic fibroblasts show slow progression of replicative senescence and resistance to SIPS by oxidative stress and OIS (Nogueira et al., 2008). Our results seem to contradict their findings, but this contradiction is possibly due to the temporal activation of Akt in the post-treatment period. The Akt-mediated pathway, which promotes cell survival and proliferation, might be beneficial for cells recovering proteostasis. These results, taken together, indicate that it is likely that mTOR inhibition with rapamycin attenuates MG132- or BAFA1-induced SIPS by repressing intracellular and mitochondrial ROS levels, restoring proteostasis, and promoting cell proliferation.

In conclusion, we have shown the temporal depletion of antioxidant proteins SOD2 and GPx4 in mitochondria, excess production of mitochondrial ROS, changes in mitochondrial function, and PGC-1α-mediated mitochondrial biogenesis in cells with impaired proteostasis. Hence, excessive ROS, possibly generated from increased mass of mitochondria, can cause nuclear DNA damage, cell cycle arrest, and eventual cellular senescence. Our study proposes a possible pathway from the disturbance of proteostasis to cellular senescence via functional changes in mitochondria.

## 4 EXPERIMENTAL PROCEDURES

### 4.1 Cell culture

MRC-5 cells (30 PDL) were obtained from the JCRB Cell Bank. Cells were maintained in EMEM (Fujifilm Wako Pure Chemical Corp, Osaka, Japan) supplemented with 10% fetal bovine serum (Biological Industries, Cromwell, CT) and penicillin-streptomycin (Nacalai Tesque, Kyoto, Japan) at 37°C in a humidified atmosphere of 5% CO_2_. Cells were passaged every 2 days.

### 4.2 MG132 and BAFA1 treatment

For MG132 or BAFA1 treatment, we plated 1 × 10^4^ cells a well in a 96-well clear bottom black plate, allowed them to grow for 16-24 h, and then changed the culture medium to EMEM containing MG132 (ChemScene, Monmouth Junction, NJ) or BAFA1 (Sigma-Aldrich). As a pilot experiment, we tested the viability of cells treated with MG132 (0.2-1 μM) or BAFA1 (8-15 nM) at several different concentrations to determine the appropriate concentration of each drug. On day 5 of drug treatment, the culture medium was replaced with regular EMEM that contained no drug. For rapamycin co-treatment, cells were treated with 10 nM rapamycin (LC Laboratories, Woburn, MA).

### 4.3 Immunoblotting and antibodies

Cells cultured in 6 cm dishes were trypsinized and harvested. Cell pellets were lysed in RIPA buffer (20 mM Tris-HCl, pH7.4, 100 mM NaCl, 0.5% Nonidet P-40, 0.05% SDS, 1 mM EDTA, protease inhibitors). Protein concentration was determined using Quick Start Bradford dye (Bio-Rad). Whole cell extracts (10-15 μg) were mixed with 5×SDS sample buffer containing 2-mercaptoethanol, heated at 95°C for 3 min, and then loaded onto an SDS-polyacrylamide gel (SDS-PAGE). For the immunoblot analysis of fixed SOD2 complex, cells were fixed in 4% paraformaldehyde (PFA) in phosphate-buffered saline (PBS) for 10 min at 25°C, washed with PBS, lysed in RIPAP buffer, sonicated briefly, and centrifuged at 21,500 ×g for 5 min. The supernatant was mixed with 5×SDS sample buffer, incubated at 95°C for 3 min or 37°C for 15 min, and loaded onto SDS-PAGE. Proteins were transferred to a PVDF membrane and incubated in 1% Western blocking reagent (Sigma-Aldrich) at 25°C for 1 h. The membrane was then incubated overnight with antibodies diluted in 0.5% Western blocking reagent. Antibodies used in this study are summarized in a supplemental file. The chemiluminescence signal was visualized with ECL prime reagent (GE Healthcare) or ImmunoStar LD (Fujifilm), and detected with LAS4000 (GE Healthcare). The intensity of the obtained protein band was quantified using ImageJ software ver. 2.0.0.

### 4.4 SA-β-gal staining and senescence assays

SA-β-gal staining was performed as described previously (Dimri et al., 1995). Quantitation of SA-β-gal activity was evaluated using Cellular Senescence Plate Assay Kit - SPiDER-βGal, a fluorogenic substrate for β-galactosidase (Dojindo, Kumamoto, Japan), following the manufacturer’s instructions.

### 4.5 Immunofluorescence

For immunofluorescence analyses, cells were fixed in 4% PFA/PBS or methanol, permeabilized with 0.5% Triton X-100/PBS for 10 min, incubated in Blocking One Histo (Nacalai Tesque, Kyoto, Japan) for 1 h, and incubated with diluted primary antibody overnight. Antibodies used in this study are summarized in a supplemental file. After several PBS washes, cells were incubated with a secondary antibody and stained with Hoechst 33342. Cells were observed under a fluorescence microscope IX83 or a laser scanning microscope FV1200 (Olympus, Tokyo, Japan).

### 4.6 Measurement of ROS and NO

Production of hydroxyl radical (.OH) and hypochlorous acid (HClO) were measured using the OxiORANGE reagent (Goryo Chemical, Sapporo, Japan). H_2_O_2_ and NO were detected by HYDROP and diaminofluorescein-FM diacetate (DAF-FM DA)(Goryo Chemical, Sapporo, Japan), respectively. Cells plated in a 96-well Black IsoPlate (PerkinElmer, Waltham, MA) were incubated with 0.5 μM OxiORANGE, 1 μM HYDROP, or 1 μM DAF-FM DA diluted in EMEM containing FBS for 30 min at 37°C, and then washed with 100 μL HBSS once. Fluorescence (Ex/Em = 535/595 nm for OxiORANGE, 485/535 nm for HYDROP and DAF-FM DA) was measured by using FilterMax F5 (Molecular Devices, San Jose, CA). Normalized values were obtained by dividing the fluorescent intensities of HYDROP and DAF-FM DA by protein concentration or dividing the fluorescent intensities of OxiORANGE by those of nuclei stained with Hoechst 33342 (Dojindo, Kumamoto, Japan).

### 4.7 Staining of protein aggregation

Protein aggregation was visualized using ProteoStat Aggresome Detection Reagent (Enzo Life Science, Farmingdale, NY), according to the manufacturer’s protocol. For the quantitative analysis, cells plated in 48-well black plates were fixed with 4% PFA/PBS, permeabilized with 0.5% Triton X-100, and stained with ProteoStat Aggresome Detection Reagent for 30 min at 25°C, and observed under an all-in-one fluorescence microscope BZ-9000 (Keyence, Osaka, Japan). Fluorescence images were analyzed and quantified using ImageJ software ver. 2.0.0.

### 4.8 Live cell microscopy for mitochondrial analyses

To detect mitochondrial mass and ΔΨm, live cells were incubated with 0.5 μM MitoTracker Green FM (Thermo Fisher Scientific) and 0.5 μM MitoTracker Red CM-H2Xros (Thermo Fisher Scientific) with Hoechst 33342 diluted in EMEM containing FBS for 30 min at 37°C, and then washed with 100 μL Hank’s balanced salt solution (HBSS) once. For the detection of mitochondrial ROS, cells were incubated with 5 μM MitoSOX Red mitochondrial superoxide indicator (Thermo Fisher Scientific) in HBSS for 10 min at 37°C, washed with HBSS 3 times, and cultured in regular medium for 24 h. On the next day, cells were stained with Hoechst 33342, washed with 100 μL HBSS once, and observed under an all-in-one fluorescence microscope BZ-9000 (Keyence, Osaka, Japan). Fluorescence images were analyzed and quantified using ImageJ software ver. 2.0.0.

### 4.9 Intracellular ATP assay

Intracellular ATP levels were measured using “Cell” ATP Assay reagent (Toyo B-Net, Tokyo, Japan). The ATP level was normalized by the DNA content in cell lysates, which was quantified by SYBR Gold stain (Thermo Fisher Scientific).

### 4.10 Preparation of mitochondrial fraction

The procedure for isolation of the mitochondrial fraction from cultured cells was as described by Clayton *et al* (Clayton & Shadel, 2014) with slight modifications. The buffer volume, number of strokes, homogenizer, and pestle were optimized for 0.5-1.0 × 10^6^ cells. Frozen cell pellets were resuspended in hypotonic buffer and homogenized using a disposable plastic pestle (As One Corp., Osaka, Japan) with matched Safe-Lock tubes (Eppendorf). The detailed procedure is summarized in Fig. S6A-C.

### 4.11 Expression of SOD2-V5 protein and immunoprecipitation

The full-length of coding region of human *SOD2* was amplified by PCR with gene-specific primers, and cloned into pcDNA3.1D/V5-His-TOPO (Thermo Fisher Scientific). MRC-5 cells were transfected with the SOD2-V5 expression construct using Polyethylenimine “MAX” (Polysciences Inc.), cultured for 16 h, and then treated with MG132 or BAFA1 for 24 h. SOD2-V5 protein in 125 or 250 μg lysate was immunoprecipitated with anti-V5 antibody and Dynabeads Protein G (Thermo Fisher Scientific), and then immunoblotted with 25 μg input samples.

### 4.12 Quantitative PCR

See Supporting Information.

### 4.13 Statistical analyses

We conducted one-way ANOVA test followed by Tukey’s multiple comparison test. Two groups were compared using a non-parametric Mann-Whitney *U* test. Differences between groups with a *p* value of <0.05 were considered significant. All data were analyzed using the GraphPad Prism 5.0 software (GraphPad Software, San Diego, California USA, www.graphpad.com) and presented as the mean (SD) of the obtained values.

## ACKNOWLEDGEMENTS

We are grateful to Santa Cruz Biotechnology, Inc. (Dallas, TX) and Cosmo Bio Co., Ltd. (Tokyo, Japan) for providing the sample antibodies listed in the Supporting Information. We would like to thank Editage (www.editage.jp) for English language editing.

## AUTHOR CONTRIBUTIONS

Y. T. and Y. K. designed the research and drafted the paper; Y. T. carried out experiments and analyzed data; and I. I., T. N., and M. I. supported the analysis of oxidative stress.

## FUNDING INFO

This work was supported by a Nippon Medical School Grant-in-Aid for Medical Research 2019 and 2020.

---

**Figure S1.**
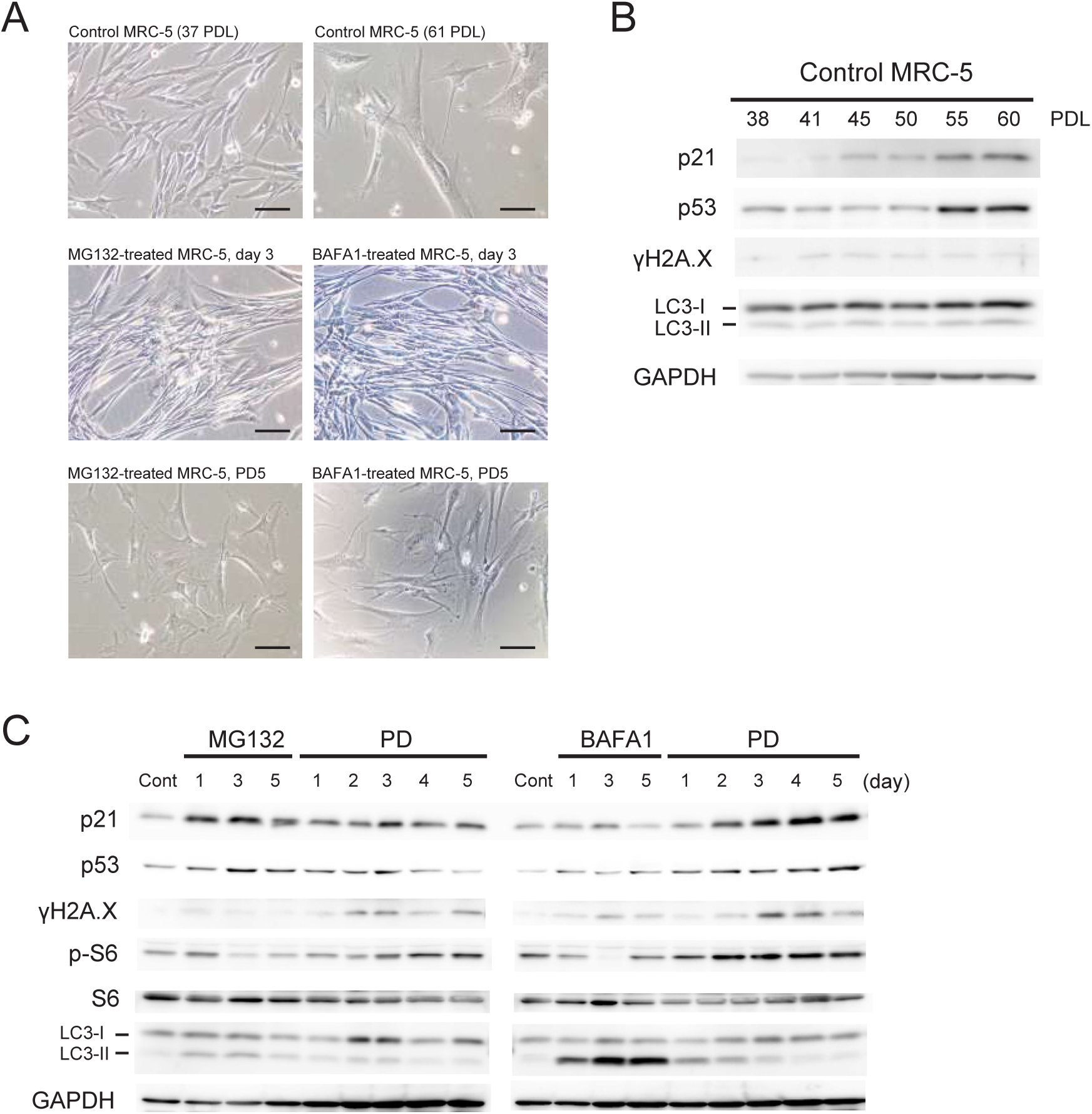
(A) Representative pictures of senescence-induced MRC-5 cells by 5-days treatment of MG132 or BAFA1 (day 3 and PD5), young (37 PDL) and aged (61 PDL) control cells. Scale bar = 100 μm. (B) Immunoblot analyses of untreated control MRC-5 cells undergoing replicative senescence (from 38 to 60 PDL) using antibodies against p21, p53, γH2A.X, LC3, and GAPDH. (C) Immunoblot analyses of MG132- or BAFA1-treated MRC-5 cells. To verify the reproducibility of the protein expressions shown in Fig. 1g, we repeated same immunoblot analyses using differently prepared whole cellular samples.

**Figure S2.**
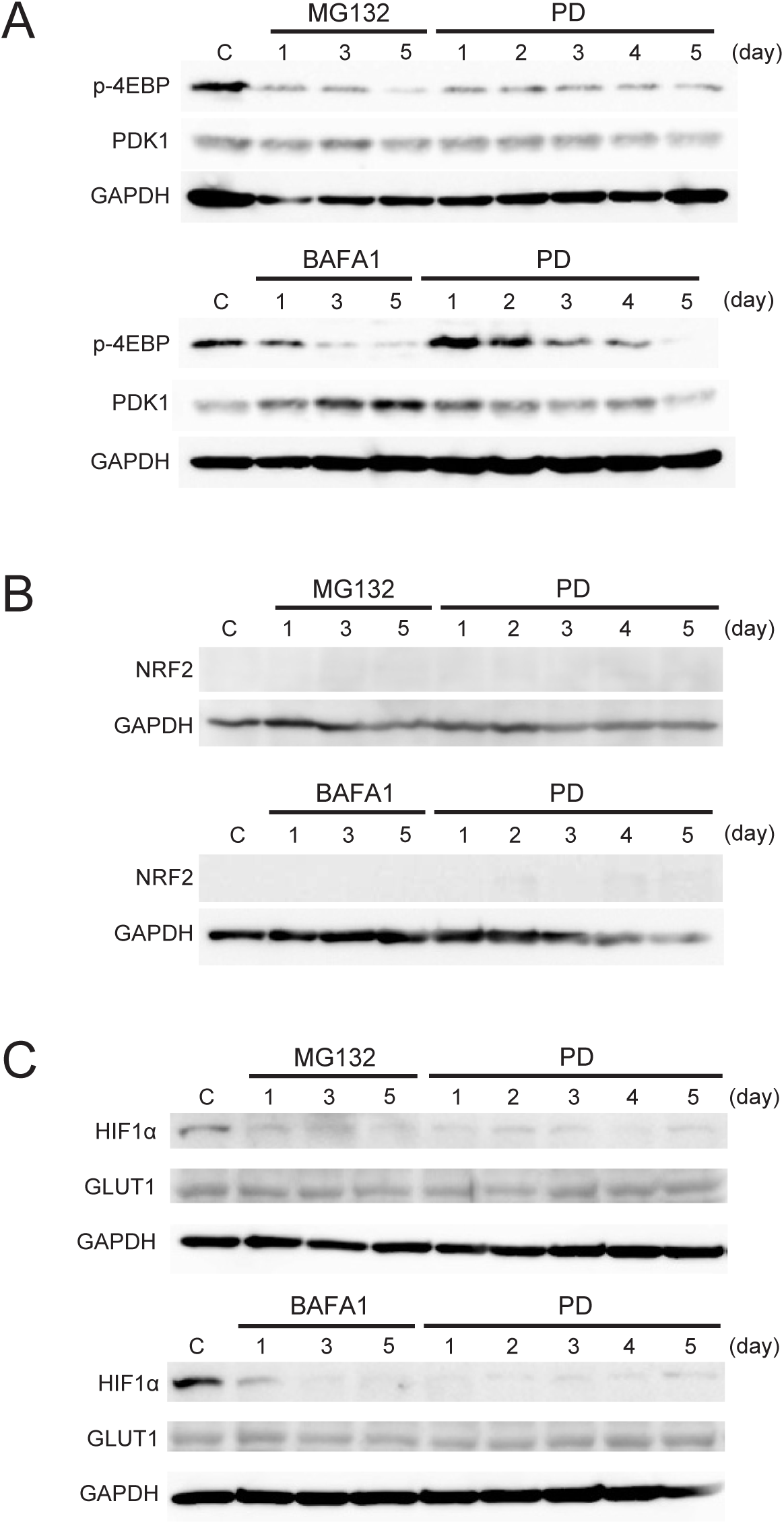
(A) Immunoblot analyses of MG132- or BAFA1-treated cells using antibodies against p-4EBP, PDK1, and GAPDH. C, untreated control cells. (B) Immunoblot analyses of MG132- or BAFA1-treated cells using antibodies against NRF2 and GAPDH. (C) Immunoblot analyses of MG132- or BAFA1-treated cells using antibodies against HIF1α, GLUT1, and GAPDH.

**Figure S3.**
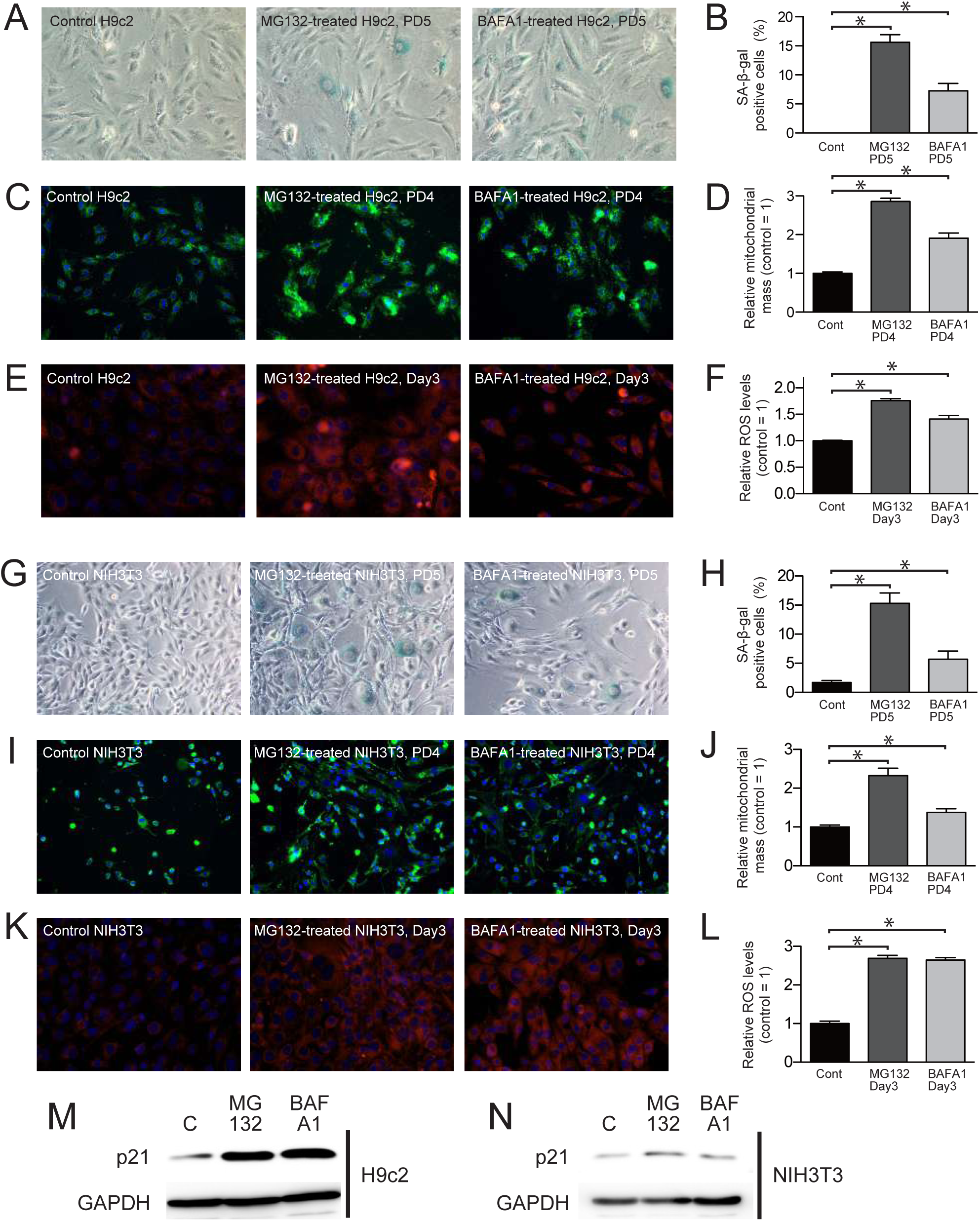
Premature senescence of H9c2 rat cardiac myoblasts (A-F) and NIH3T3 mouse fibroblasts (G-L) induced by MG132- or BAFA1-treatment. After 5-days treatment, percentages of SA-β-gal positive cells (A, B, G, and H), mitochondrial mass evaluated using MitoTracker Green FM (C, D, I, and J), and intracellular ROS levels evaluated using OxiORANGE (E, F, K, and L) significantly increased in both cell lines. Upregulation of p21 was also examined by the immunoblot analyses of H9c2 (M) and NIH3T3 (N) on PD5. Values are shown as the mean (SD) from four measurements. The asterisk (*) indicates *p* < 0.05 by a non-parametric Mann-Whitney *U* test for control vs. treated cells.

**Figure S4.**
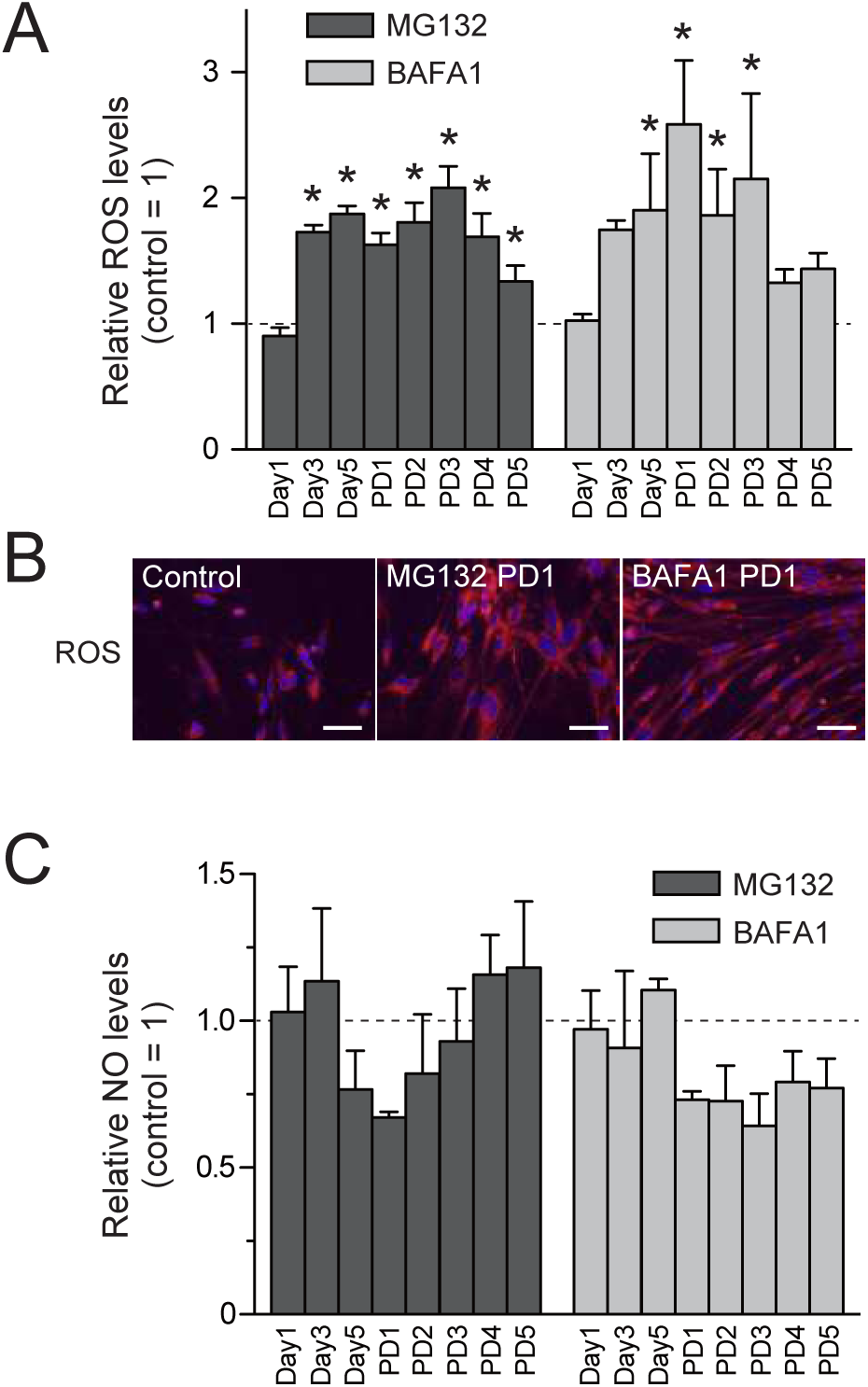
Prolonged disturbance of proteostasis by MG132 or BAFA1 enhances ROS but not NO production. (A) Quantitation of relative intracellular ROS levels using OxiORANGE reagent that selectively detects hydroxyl radical (.OH) and hypochlorous acid (HClO). The fluorescence intensity for OxiORANGE (Ex/Em = 535/595) was normalized by Hoechst 33342 intensity (Ex/Em = 340/465). Values are shown as the mean (SD) from four wells of a 96-well plate. The asterisk (*) indicates *p* < 0.05 by one-way ANOVA, followed by Tukey’ s multiple comparison test for control vs. treated cells. (B) Representative images of control and MG132- or BAFA1-treated (PD1) MRC-5 cells stained for the OxiORANGE (red) and Hoechst 33342 (blue). Scale bar = 50 μm. (C) Quantitation of relative intracellular ΝΟ levels by using DAF-FM DA fluorescent dye that selectively detects ΝΟ. The fluorescence intensity for DAF-FM DA was normalized by total protein concentration. Values are shown as the mean (SD) from four wells of a 96-well plate.

**Figure S5.**
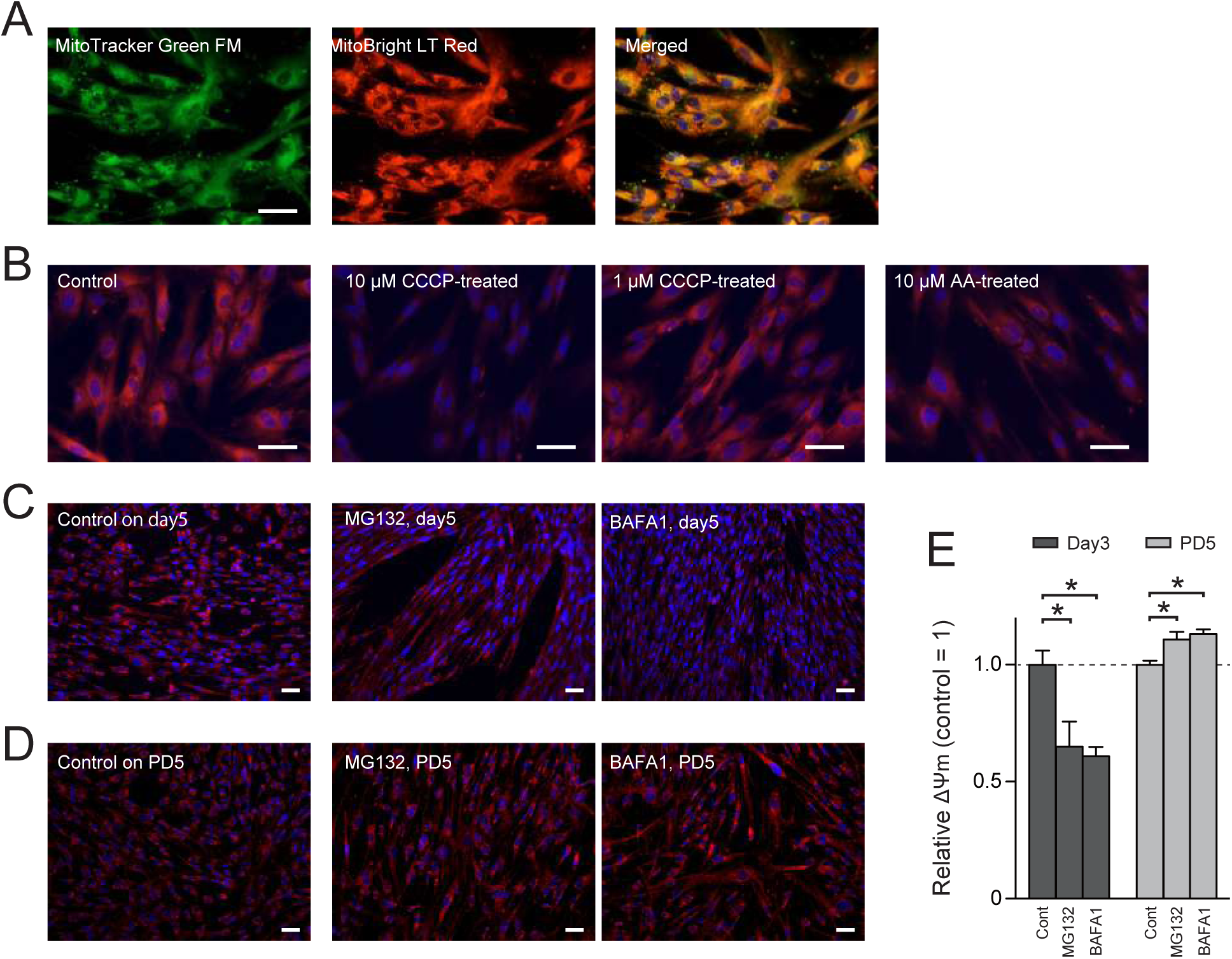
Evaluation of mitochondrial membrane potential (ΔΨm) using MitoBright LT Red (MBR). (A) Double staining of MRC-5 cells using MitoTracker Green FM and MBR (Dojindo, Kumamoto, Japan). Both reagents stained mitochondria in a similar manner. (B) MBR fluoresces in a ΔΨm-dependent manner. MBR-stained MRC-5 cells were treated with the mitochondria uncoupler carbonyl cyanide m-chlorophenylhydrazone (CCCP) or the respiration inhibitor antimycin A (AA) at the indicated concentrations for 30 min. MBR signals were markedly reduced by these treatments. (C and D) MRC-5 cells treated with MG132 or BAFA1 were stained with MBR on day 5 (C) or PD5 (D) to evaluate the ΔΨm. (E) Relative ΔΨm of MG132- or BAFA1-treated MRC-5 cells on day 3 or PD5. MG132- or BAFA1-treated MRC-5 cells were co-stained with MitoTracker Green FM and MBR and first segmented with the MitoTracker Green FM signal using Image J, and then their area was calculated. Next, the signal of MBR within the MitoTracker Green FM area was measured. Finally, the MBR signal was divided by the area of MitoTracker Green FM to obtain the ΔΨ m signal per unit area. This was done to complement the results in Fig. 3b. Similar to the results in Fig. 3b, relative ΔΨm decreased during MG132 or BAFA1 treatment, but increased after the treatment. Values are shown as the mean (SD) from six measurements. The asterisk (*) indicates *p* < 0.05 by a non-parametric Mann-Whitney *U* test for control vs. treated cells. Scale bar = 20 μm.

**Figure S6.**
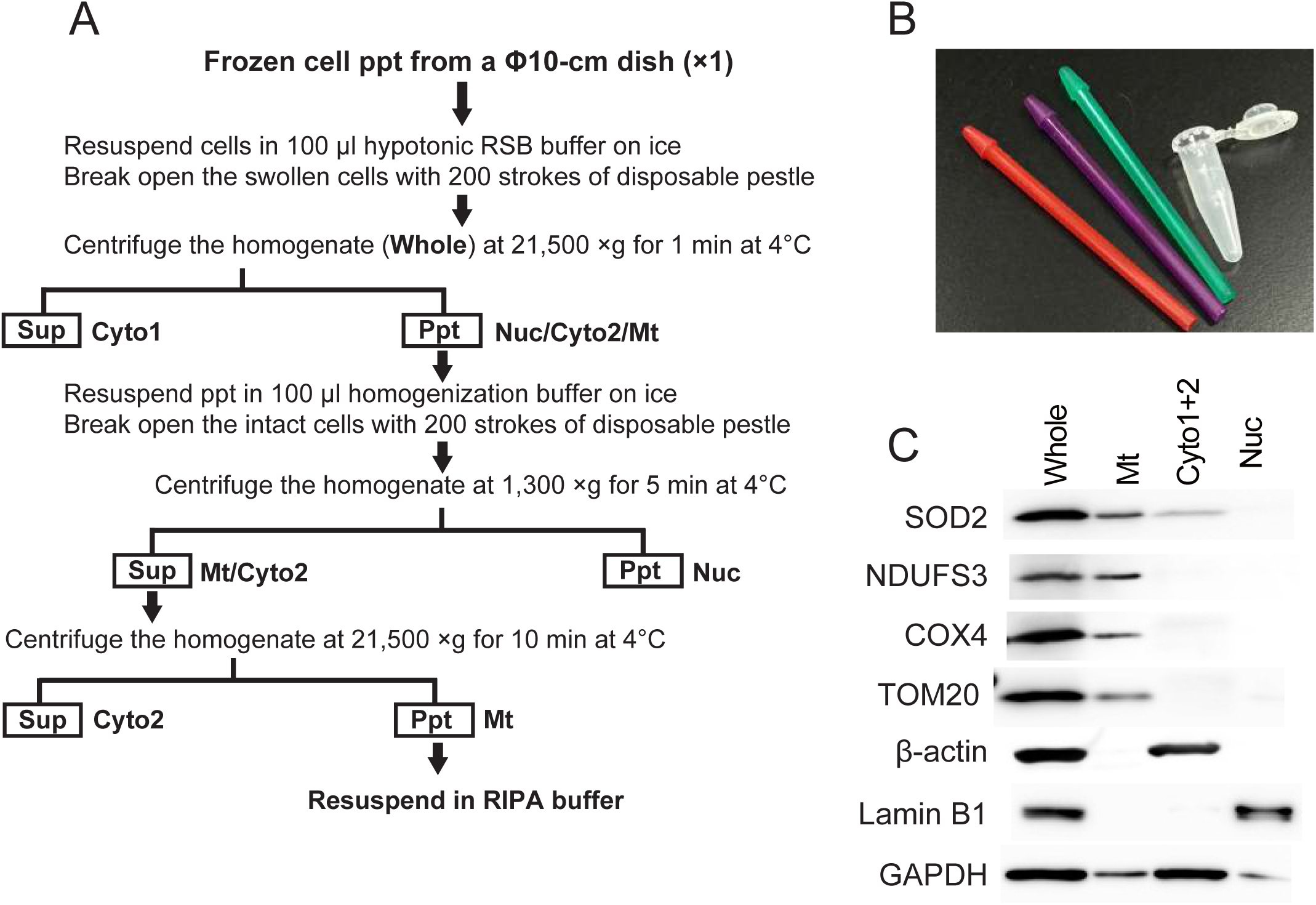
Preparation of crude mitochondrial fraction from MRC-5 cells. (Α) A flow chart for preparation of mitochondrial fraction from frozen cell pellet at a microscale. Cells were homogenized in hypotonic RSB buffer, then unbroken cells were repeatedly homogenized in a homogenization buffer. For the immunoblot analyses, solubilized mitochondrial proteins in RIPA buffer containing 0.5% NP40 (Mt) were applied. Hypotonic RSB buffer, 10 mM Tris-HCl, pH7.4, 10 mM NaCl, 1.5 mM MgCl2, protease inhibiter; Homogenization buffer, 5 mM Tris-HCl, pH7.4, 1.5 mMgCl2, 250 mM sucrose, protease inhibitor; RIPA buffer, 20 mM Tris-HCl, pH7.4, 100 mM NaCl, 0.5% Nonidet P-40, 0.05% SDS, 1 mM EDTA, protease inhibitors; Cyto, cytoplasmic fraction; Nuc, nuclear fraction; Whole, whole cellular extract. (B) Disposable plastic pestles and a matched Safe-Lock tube used in this method. (C) Immunoblot analyses of whole cellular extract (Whole), crude mitochondrial fraction (Mt), and cytoplasmic fraction (Cyto1 and 2) prepared from untreated control MRC5 cells using antibodies against SOD2, NDUFS3, COX4, TOM20, β-actin, Lamin B1, and GAPDH.

**Figure S7.**
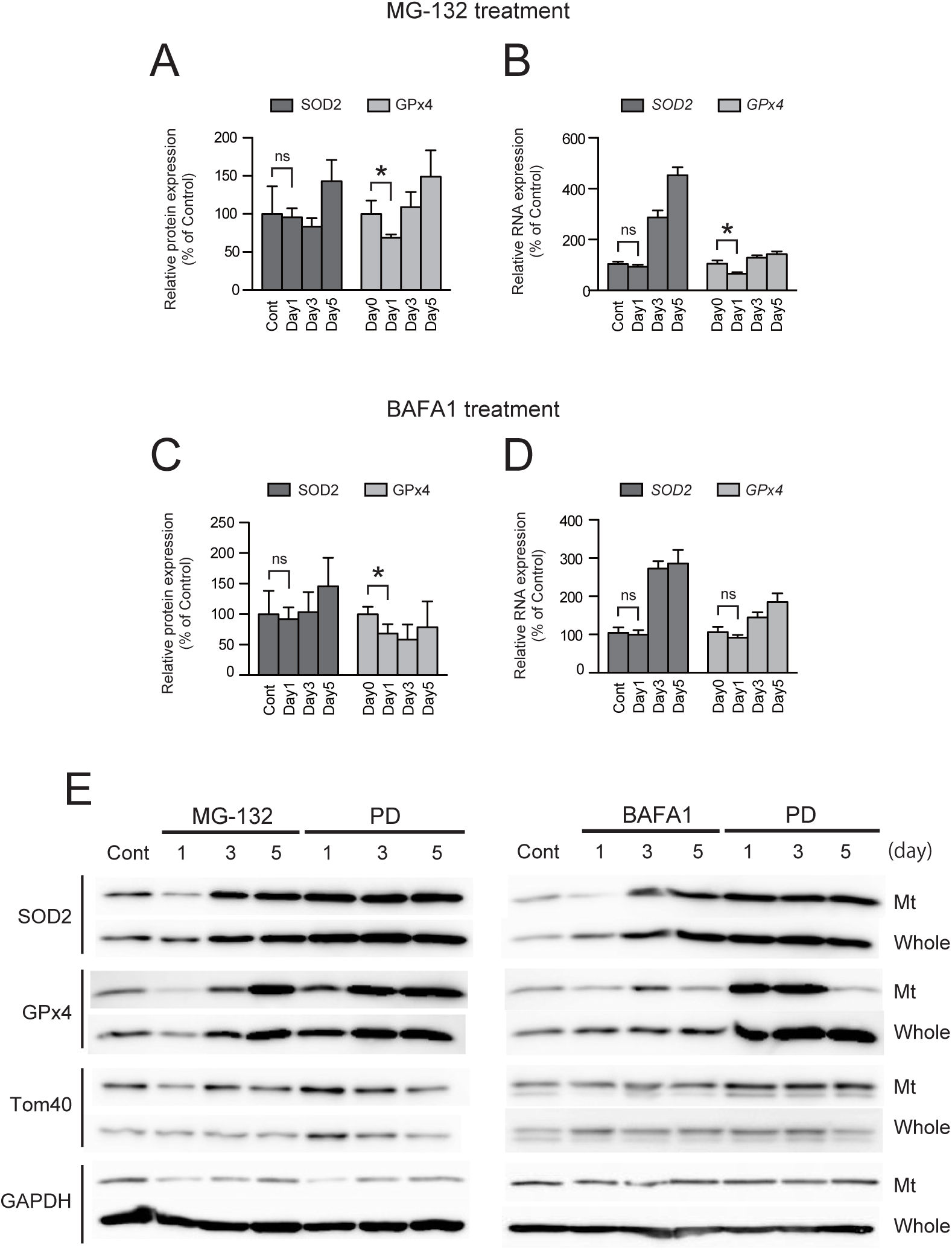
Protein and mRNA expressions of SOD2 and GPx4 in MRC-5 cells treated with MG132 (A, B, and E) or BAFA1 (C, D, and F). (A and B) Protein levels of SOD2 and GPx4 were evaluated by densitometry of each band obtained by immunoblot analyses of 4 individually prepared whole cellular lysates. Intensities of SOD2 and GPx4 bands were normalized to the intensity of GAPDH. Values are shown as the mean ± SD. The asterisk (*) indicates *p* < 0.05 by a non-parametric Mann-Whitney *U* test for control vs. treated cells. ns, not significant. (B and D) mRNA levels of *SOD2* and *GPx4* were evaluated by quantitative RT-PCR of four individual cDNAs prepared from the total RNA. (E) Immunoblot analyses of MRC-5 cells treated with MG132 (left) or BAFA1 (right). To verify the reproducibility of the protein expressions shown in Fig. 4a and 4b, we repeated immunoblot analyses for differently prepared mitochondrial and whole cellular fractions.

**Figure S8.**
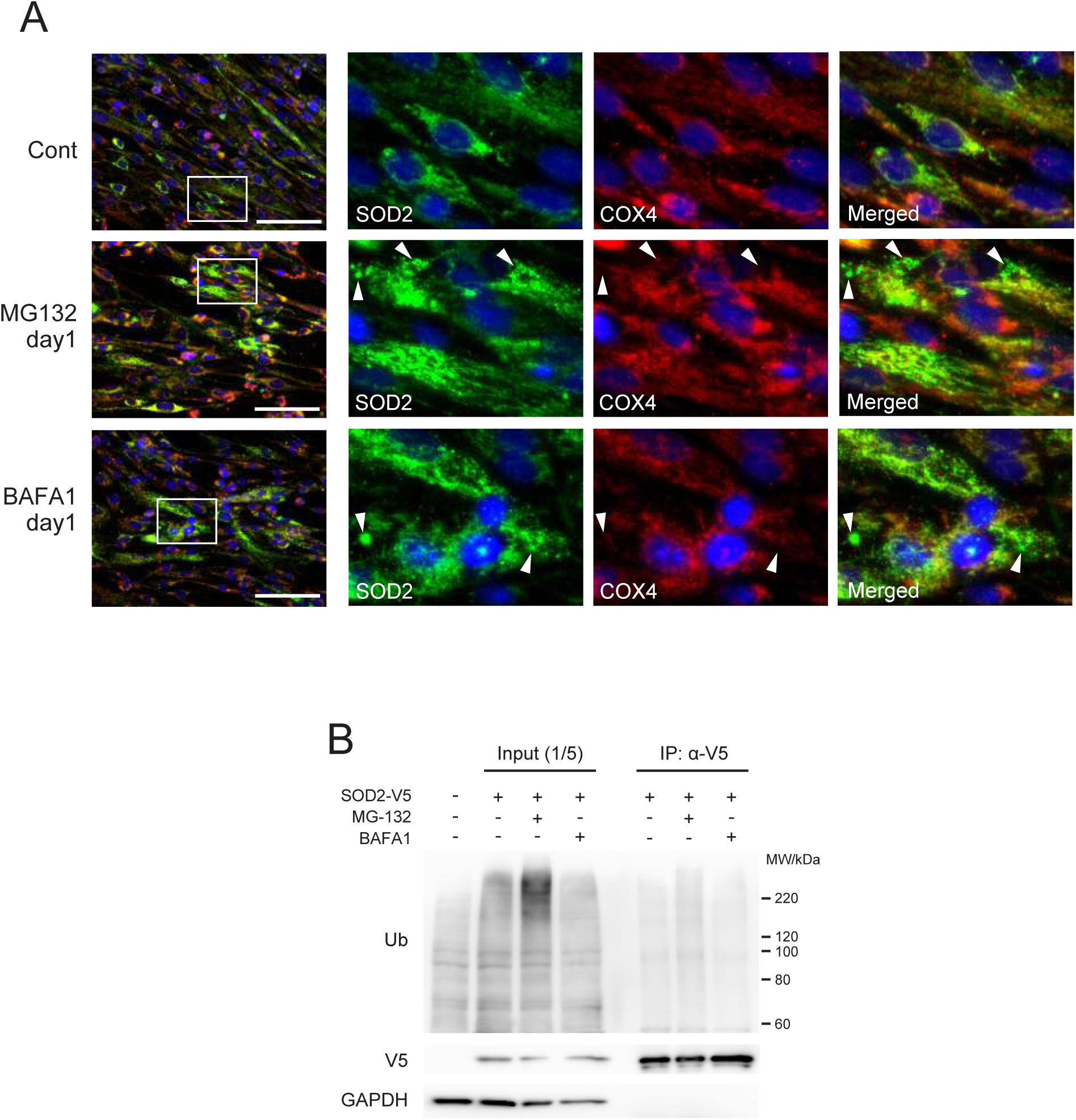
(A) Extra-mitochondrial localization of SOD2 in MG132- or BAFA1-treated MRC-5 cells on day1. Cells were co-stained with anti-SOD2 (green) and anti-COX4 (red) antibodies. Nuclei were stained with Hoechst 33342 (blue). Arrowheads indicate extra-mitochondrially localized SOD2. Immunostained cells were observed under an all-in-one fluorescence microscope BZ-9000 (Keyence). Cont, control cells. Scale bar = 100 μm. (B) SOD2 is not polyubiquitinated in MG132- or BAFA1-treated cells on day 1. SOD2-V5 was immunoprecipitated (IP) using anti-V5 antibody (α-V5), and the precipitants were subjected to immunoblot analysis with anti-ubiquitin (Ub), V5, and GAPDH antibodies.

**Figure S9.**
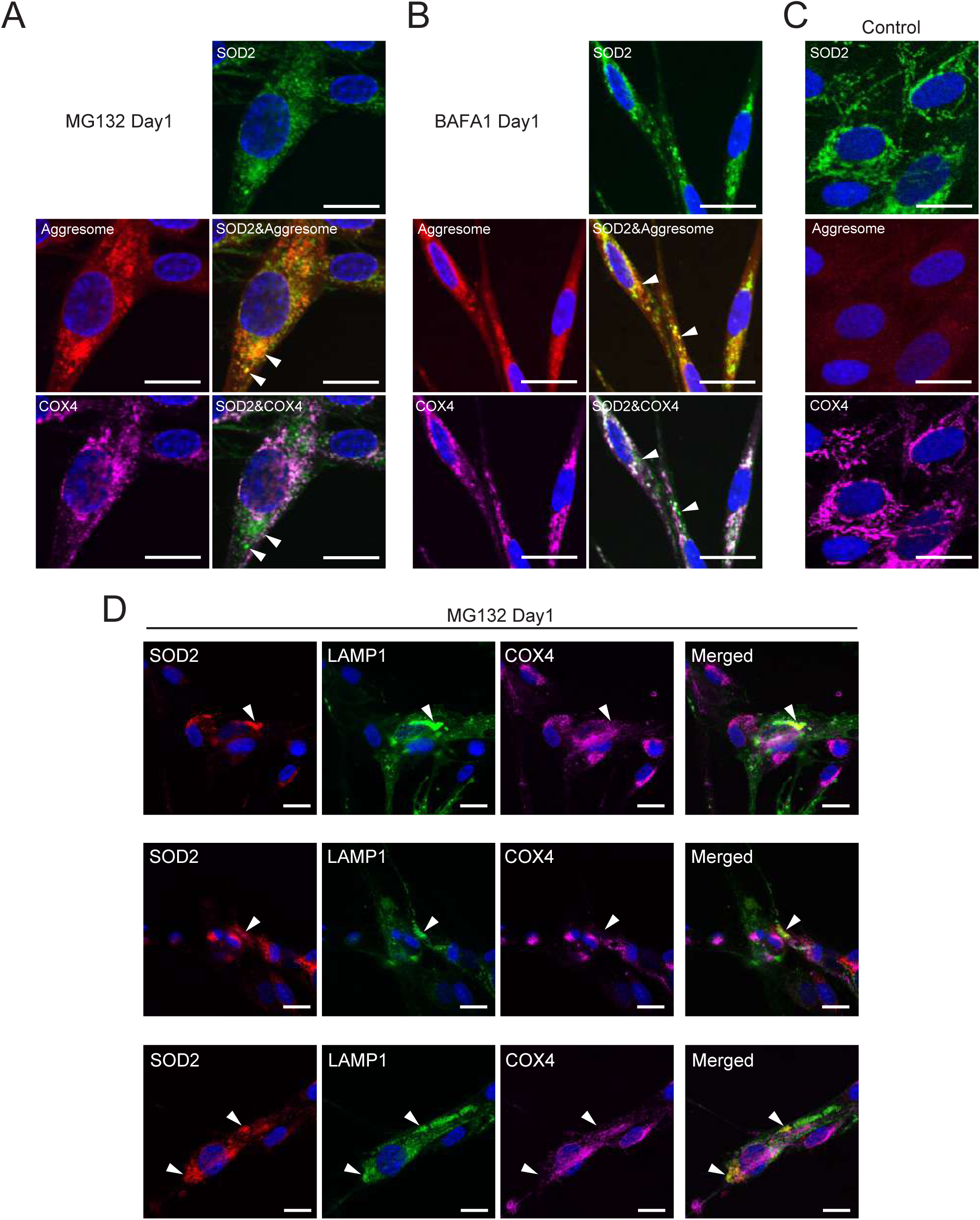
Extra-mitochondrially localized SOD2 colocalizes with aggresome. (A-C) Control, MG132- or BAFA1-treated MRC-5 cells on day 1 were co-stained with anti-SOD2 (green) and anti-COX4 (purple) antibodies. Aggresome (red) and nuclei (blue) were stained with the ProteoStat Aggresome Detection kit and Hoechst 33342, respectively. Arrowheads indicate colocalization of extra-mitochondrial SOD2 with the aggresome. (D) Extra-mitochondrially localized SOD2 colocalized with lysosomes. MRC-5 cells were transiently transfected with CellLight^TM^ Lysosomes-GFP baculovirus expressing the lysosomal associated membrane protein LAMP1-GFP fusion protein, and then treated with MG132 for one day. Cells were co-stained with anti-SOD2 (red) and anti-COX4 (purple) antibodies. Arrowheads indicate colocalization of extra-mitochondrial SOD2 with LAMP1. Immunostained cells were observed under a laser scanning microscope FV1200 (Olympus). Scale bar = 20 μm.

**Figure S10.**
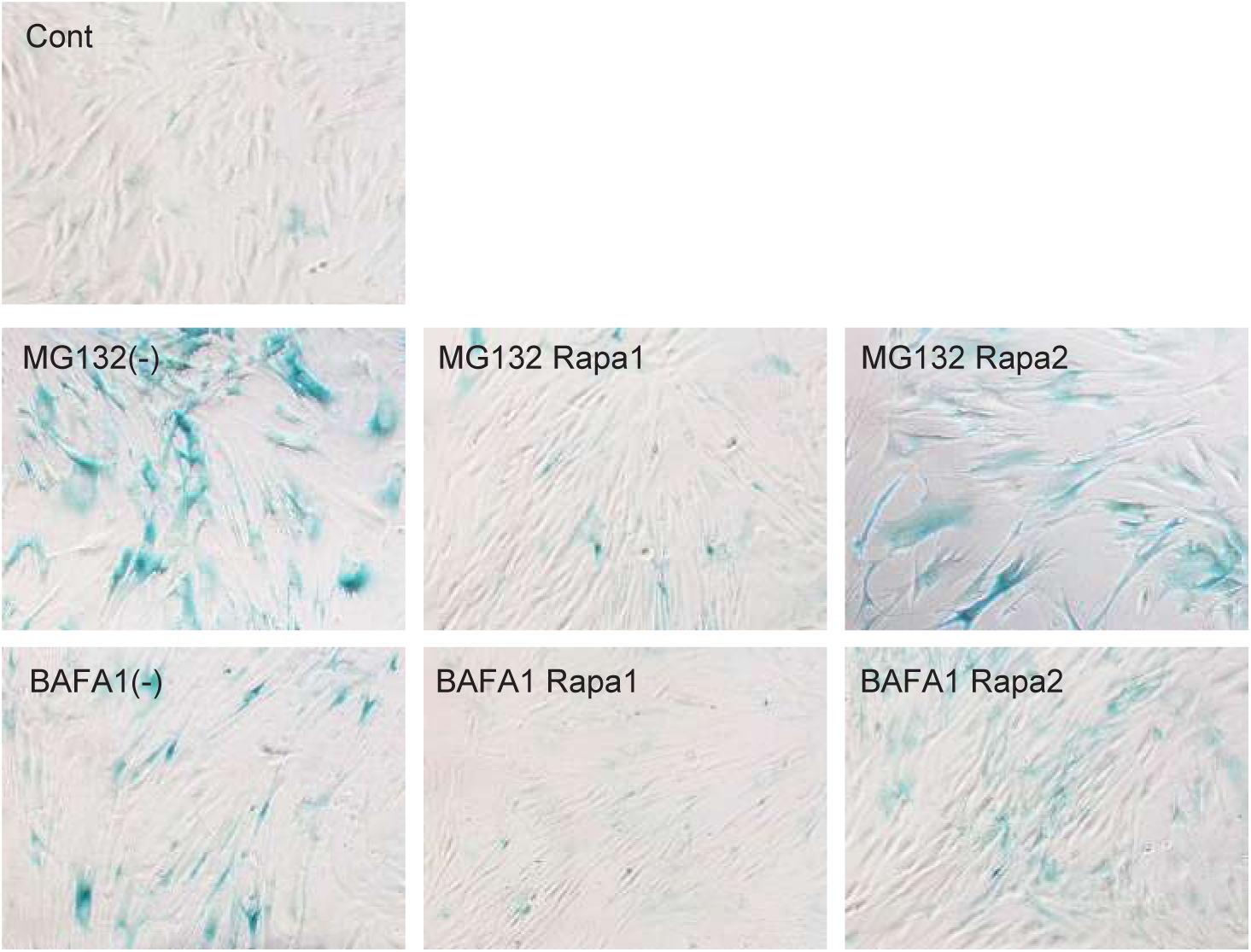
Representative pictures of control and MG132- or BAFA1-treated MRC-5 cells stained with SA-β-gal. Cells were co-treated with rapamycin and either MG132 or BAFA1 for 5 days, respectively, and stained on PD5 (cytoplasmic blue precipitate). Control, untreated young (39 PDL) MRC-5 cells. (-), cells treated with only MG132 or BAFA1; Rapa1 and Rapa2, cells co-treated with rapamycin and either MG132 or BAFA1 as shown in Fig. 6a.

**Figure S11.**
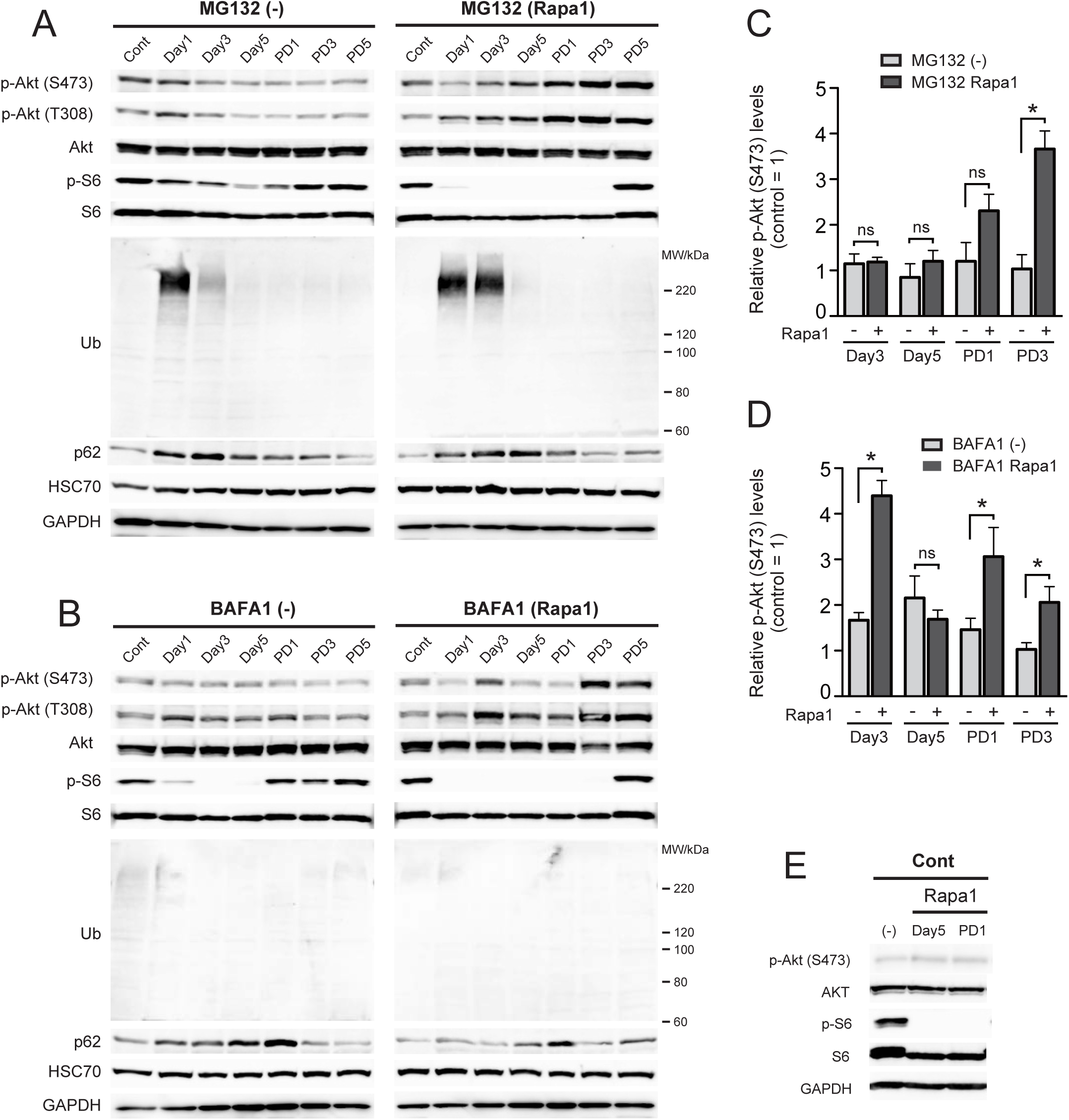
Co-treatment of rapamycin with MG132 or BAFA1 enhances the Akt phosphorylation and the ubiquitination in MRC-5 cells. (A) Immunoblot analyses of MG132-treated cells (-), MG132 and rapamycin co-treated cells (Rapa1). (B) Immunoblot analyses of BAFA1-treated cells (-), BAFA1 and rapamycin co-treated cells (Rapa1). (C) Quantitative analyses of the Akt phosphorylation of MG132 (-)- or MG132 and rapamycin (Rapa1)-treated cells. (D) Quantitative analyses of the Akt phosphorylation of BAFA1 (-)- or BAFA1 and rapamycin (Rapa1)-treated cells. Phosphorylated Akt (p-Akt) at Ser473 were evaluated by densitometry of each band obtained by immunoblot analyses of four individually prepared whole cellular lysates. Values are shown as the mean (SD). The asterisk (*) indicates *p* < 0.05 by a non-parametric Mann-Whitney *U* test for MG132 or BAFA1 single-treated cells vs. rapamycin co-treated cells. ns, not significant. (E) Immunoblot analyses of MRC-5 cells treated with only rapamycin for 5 days. (-), cells with no rapamycin treatment.

**Figure S12.**
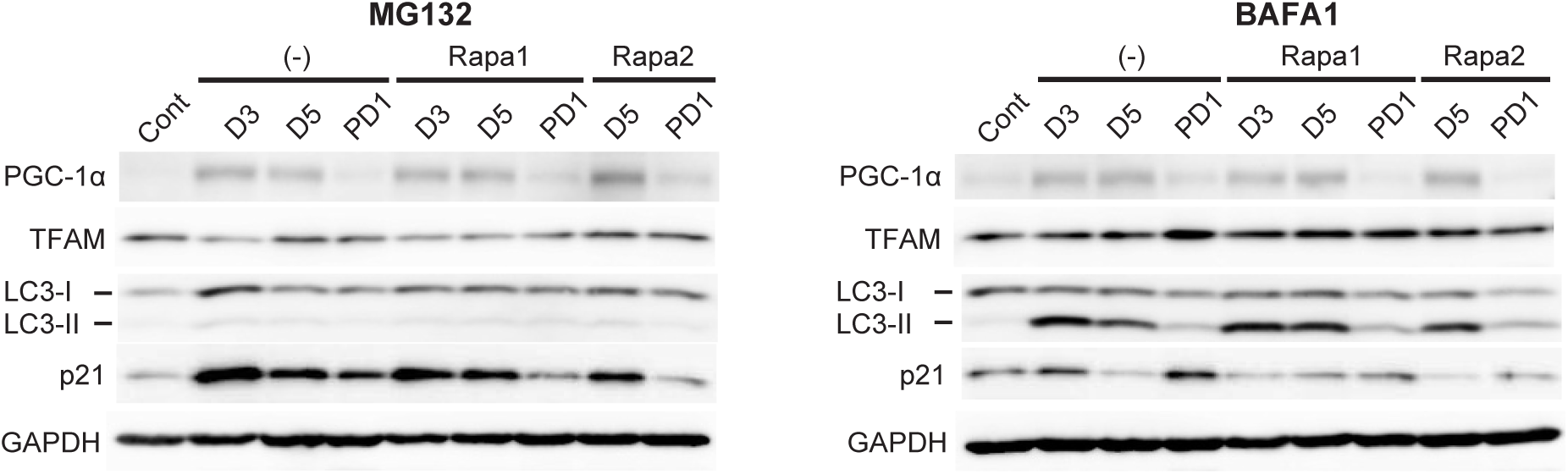
Immunoblot analyses of MG132 (left)- or BAFA1 (right)-treated MRC-5 cells with (Rapa1 and Rapa2) or without (-) rapamycin at day 3 (D3), day 5 (D5), and PD1.

## DATA AVAILABILITY STATEMENT

The data that support the findings of this study are available from the corresponding author upon reasonable request.

## Supporting information

### 1. Antibodies used in this study

#### 1st antibodies

anti-p21 (SX118, sc-53870, Santa Cruz Biotechnology, mouse monoclonal)

anti-S6 (C-8, sc-74459, Santa Cruz Biotechnology, mouse monoclonal)

anti-p-S6 (50.Ser 235/236, sc-293144, Santa Cruz Biotechnology, mouse monoclonal)

anti-p53 (DO-1, sc-126, Santa Cruz Biotechnology, mouse monoclonal)

anti-p-Histone H2A.X (Ser 139, sc-517348, Santa Cruz Biotechnology, mouse monoclonal)

anti-mtTFA (TFAM)(C-9, sc-376672, Santa Cruz Biotechnology, mouse monoclonal)

anti-PGC-1α (D-5, sc-518025, Santa Cruz Biotechnology, mouse monoclonal)

anti-Parkin (PRK8, sc-32282, Santa Cruz Biotechnology, mouse monoclonal)

anti-SOD2 (A-2, sc-133134, Santa Cruz Biotechnology, mouse monoclonal)

anti-GPx-4 (E-12, sc-166570, Santa Cruz Biotechnology, mouse monoclonal)

anti-NDUFS3 (D-4, sc-374282, Santa Cruz Biotechnology, mouse monoclonal)

anti-Tom20 (F-10, sc-17764, Santa Cruz Biotechnology, mouse monoclonal)

anti-Tom40 (D-2, sc-365467, Santa Cruz Biotechnology, mouse monoclonal)

anti-NRF1 (G-5, sc-515360, Santa Cruz Biotechnology, mouse monoclonal)

anti-NRF2 (M200-3, Medical & Biological Laboratories Co., ltd., Japan, mouse monoclonal)

anti-Cox-4 (#3638-100, BioVision, rabbit polyclonal) for immunoblot

anti-COX4 (PM063MS, MBL Life Science, rabbit polyclonal) for immunostaining

anti-LC3 (PM036, MBL Life Science, rabbit polyclonal)

anti-GAPDH (G8795, Sigma-aldrich, mouse monoclonal)

anti-p-AMPKα (#2535, Cell Signaling Technology)

anti-AMPKα (#2532, Cell Signaling Technology)

anti-β-actin (C4, sc-47778, Santa Cruz Biotechnology, mouse monoclonal)

anti-Lamin B1 (ab16048, abcam, rabbit polyclonal)

anti-HSP70 (C92F3A-5, sc-66048, Santa Cruz Biotechnology, mouse monoclonal)

anti-HSC70 (B-6, sc-7298, Santa Cruz Biotechnology, mouse monoclonal)

anti-LAMP-2 (H4B4, sc-18822, Santa Cruz Biotechnology, mouse monoclonal)

anti-AKT (#9272S, Cell Signaling Technology, rabbit polyclonal)

anti-p-AKT (S473) (#9271S, Cell Signaling Technology, rabbit polyclonal)

anti-p-AKT (T308) (#2965L, Cell Signaling Technology, rabbit monoclonal)

anti-Ub (P4D1, sc-8017, Santa Cruz Biotechnology, mouse monoclonal)

anti-SQSTM1 (p62) (D-3, sc-28359, Santa Cruz Biotechnology, mouse monoclonal)

anti-GLUT1 (H-43, sc-7903, Santa Cruz Biotechnology, rabbit polyclonal)

anti-HIF1a (NB100-123, Novus Biologicals, mouse monoclonal)

anti-p-4EBP (62.Ser 65, sc-293124, Santa Cruz Biotechnology, mouse monoclonal)

anti-PDK1 (4A11F5, sc-293160, Santa Cruz Biotechnology, mouse monoclonal)

anti-V5 (R960-25, Thermo Fisher, mouse monoclonal)

#### 2nd antibodies

HRP-conjugated goat anti-mouse IgG (A9044, Sigma-Aldrich)

HRP-conjugated anti-rabbit IgG (NA934, GE healthcare)

HRP-conjugated anti-rabbit IgG (#62-6120, Zymed Laboratories)

DyLight 488-conjugated anti-mouse IgG (#35503, Thermo Fisher, mouse monoclonal)

DyLight 594-conjugated anti-mouse IgG (#35510, Thermo Fisher, mouse monoclonal)

DyLight 649-conjugated anti-rabbit IgG (#611-143-122, Rockland Immunochemicals, Inc)

### 2. Quantitative PCR (qPCR)

#### qPCR primers

For human *SOD2*

Hs_SOD2_RT-UP1, 5’-AACGGGGACACTTACAAATTGCT-3’

Hs_SOD2_RT-LP1, 5’-CCCAGTTGATTACATTCCAAATAGCTT-3’

For human *GPx4*

Hs_GPx4_RT-UP1, 5’-CGCGGGCTACAACGTCAAATTCGAT-3’

Hs_GPx4_RT-LP1, 5’-CCACGCAGCCGTTCTTGTCGAT-3’

For 18S ribosomal RNA

LEM-m18S-F, 5’-CGGCTACCACATCCAAGGAA-3’

LEM-m18S-R, 5’-GCTGGAATTACCGCGGCT-3’

#### Procedures

Total RNA was isolated from MRC-5 cells using RNeasy Mini kit (QIAGEN), and reverse-transcribed using SuperScript III (Thermo Fisher Scientific) with random primers. To quantify human *SOD2*, *GPx4,* and 18S ribosomal RNA expression by qPCR, we used SYBR Gold (Thermo Fisher Scientific) and rTaq DNA polymerase (TOYOBO, Osaka, Japan) in an Thermal Cycler Dice, Real Time System III (TAKARA BIO inc., Shiga, Japan).

